# Higher visual areas act like domain-general filters with strong selectivity and functional specialization

**DOI:** 10.1101/2022.03.16.484578

**Authors:** Meenakshi Khosla, Leila Wehbe

## Abstract

Neuroscientific studies rely heavily on a-priori hypotheses, which can bias results toward existing theories. Here, we use a hypothesis-neutral approach to study category selectivity in higher visual cortex. Using only stimulus images and their associated fMRI activity, we constrain randomly initialized neural networks to predict voxel activity. Despite no category-level supervision, the units in the trained networks act as detectors for semantic concepts like ‘faces’ or ‘words’, providing solid empirical support for categorical selectivity. Importantly, this selectivity is maintained when training the networks without images that contain the preferred category, strongly suggesting that selectivity is not domain-specific machinery, but sensitivity to generic patterns that characterize preferred categories. The ability of the models’ representations to transfer to perceptual tasks further reveals the functional role of their selective responses. Finally, our models show selectivity only for a limited number of categories, all previously identified, suggesting that the essential categories are already known.

**Teaser:** Models trained solely to predict fMRI activity from images reveal strong category selectivity in higher visual areas, even without exposure to these categories in training.

## Introduction

Primary visual cortex studies have exploited an approach that presents model organisms with abstract stimuli like oriented edges or sine-wave gratings and examines the stimuli-evoked neuronal responses [1, 2]. This approach has revealed that neurons in the visual cortex are selective to particular aspects of our visual world, thereby informing our understanding of the selectivities in the early visual system. However, detailed characterization of neuronal responses in higher-order visual areas has been more elusive due to a circular difficulty. Understanding what an area encodes requires presenting the optimal stimulus, but the selectivity of the visual area remains hidden until the optimal pattern is presented [3].

Typically, past studies used an a priori hypothesis-driven approach. This deductive approach hypothesizes the stimulus attributes represented in a region and then designs experiments with carefully selected stimuli. It has identified several category-selective regions within the human ventral temporal cortex and macaque inferotemporal cortex, including regions responding selectively to faces [4, 5, 6], places [7, 8, 9], bodies [10, 11, 12, 6], tools [13], words [14, 15], and other categories [16]. However, this approach is not exhaustive or scalable; response properties within large swathes of the sensory cortex remain elusive.

An alternative approach uses naturalistic images and videos to drive activity in the visual cortex and then encoding models to test hypotheses about voxel-level tuning [17, 18, 19, 20]. This inductive approach decouples data collection from hypothesis testing, as multiple candidate models are tested post-hoc on the same dataset. Each model is constructed to test a given hypothesis and each hypothesis is formulated in terms of explicit quantitative functional forms, e.g. low-level gabor wavelet pyramid, motion-energy pyramid, semantic encoding models etc.

Recently, representations from deep neural networks have been used to construct encoding models for neural responses along the ventral visual pathway in humans and non-human primates [21, 22, 23, 24, 25, 26, 27, 28]. Optimizing for tasks like object recognition can lead to representations that accurately predict activity in the ventral visual pathway, and different tasks can align with specific brain regions [29]. This suggests that these artificial and biological networks could share computational goals, offering a new way to test computational hypotheses about the brain. Under this framework, the task that a network is trained on determines the hypothesis.

All the approaches above rely on specific underlying hypotheses. In the deductive approach, the hypothesis governs the selection of the narrow range of stimuli. In the inductive system identification approach, hypotheses are specified via an encoding model. When an encoding model uses representations from task-optimized deep neural networks, the hypothesis is specified by the network task. In contrast, a hypothesis-neutral modeling approach utilizes fewer a priori assumptions, so the resulting models are more impartial, flexible and effective in revealing tuning properties throughout the visual system.

One such approach is response optimization, i.e., fitting model parameters to reproduce the brain responses related to stimuli directly. When successful in predicting new responses and generalizing to unseen contexts, the response-optimization approach can facilitate the discovery of unknown neuronal tuning properties or provide strong tests for existing theories.

To date, the amount of data available to fit response-optimized models was often insufficient. However, recent advances in large-scale data collection, where the stimuli are not tethered to any hypothesis, present an opportunity to revisit this approach. Indeed, several recent studies have successfully built response-optimized models of early visual cortex [30, 31, 32, 33].

However, we still do not know how such an approach generalizes to higher-order visual areas and what it might reveal about them. Thus, we adopt a hypothesis-neutral approach and systematically characterize selectivity and functional specialization in four human higher-level visual regions of interest (ROIs) via response-optimized models. We focus on the fusiform face area (FFA) [4], the extrastriate body area (EBA) [10], the visual word form area (VWFA) [14], and the retrosplenial cortex (RSC, a visual place selective area) [34]. We emphasize that our approach is hypothesis-neutral in the sense of model construction; i.e., the models are not constructed to test any particular hypothesis, rather they are largely informed by the data. First, we leverage the unprecedented scale and quality of the massive 7T fMRI Natural Scenes Dataset (NSD) [32] to train a deep neural network model to predict activity of voxels in each ROI. Each network is randomly initialized (not pre-trained on any task) and is optimized to predict the activity of all voxels in the ROI from the stimulus image. Our models achieve high prediction performance, on par or outperforming state-of-the-art task optimized models. Then, we ask what functional properties emerge spontaneously in our response-optimized models. We examine the trained networks through structural analysis (feature visualizations), as well as functional analysis (feature verbalization) by running high-throughput experiments with these models on large-scale probe datasets and dissecting the evoked network activations [35, 36]. Strikingly, despite no category-level supervision, the units in the optimized networks act as detectors for high-level visual concepts like ‘faces’ (in the FFA model) or ‘words’ (in the VWFA model), providing one of the strongest evidences for categorical selectivity in these ROIs.

The observed strong semantic selectivity raises another important question: are the ROIs simply functioning as processing units for their preferred category (e.g., simply deciding whether the stimulus contains a face or a word) or are they a by-product of a non-category-specific visual processing mechanism? To probe this, we create selective deprivations in the visual diet of these response-optimized networks and study the selectivity of model neurons in the resulting ‘deprived’ networks. We find that the resulting models still demonstrate high selectivity for the preferred category, even in the absence of any experience with the preferred category. This suggests that the same “filters” used to encode the preferred category are used to encode other non-specific natural images as well. The results presented here further indicate that category-selective voxels do not respond to their preferred category by the unique structure of that category but perhaps by some other general structural characteristics that other stimuli may share with the preferred category.

We further demonstrate that the proposed models generalize remarkably and selectively to different perceptual tasks. Representations from the FFA model can predict facial identity while those from the RSC, VWFA and EBA models can discriminate between spatial layouts of different indoor scenes, letters and body poses respectively, revealing important functional distinctions between different ROIs.

Finally, in a retrospective analysis, we asked if this approach would work even when we do not start apriori with the ROIs, i.e., would the category-selective regions reveal themselves when we train response-optimized models on responses from a sufficiently broad set of voxels across the entire ventral visual stream, and apply single unit selectivity analysis techniques on the optimized model? Our experiments yielded an affirmative answer to this question, illustrating the success of this unbiased computational approach in revealing the representational structure of the human visual cortex. Further, this experiment also suggests that faces, bodies, text and food [37, 38, 39] are the categories that selectively activate the highest number of voxels in the ventral visual cortex and provides evidence *against* the existence of single voxels or regions selective for thousands of other visual concepts covered in our broad network interpretability dictionary beyond these categories.

Together with this new class of data-driven models for higher order visual areas and novel model interpretability techniques, our study illustrates that response-driven deep neural network models of visual cortex can serve as powerful and unbiased tools for probing the nature of representations and computations in the brain.

## Results

We trained deep neural network models directly to predict the brain activity related to viewing natural images. We diverged from the current computational neuroscience approach of extracting image representations from networks previously optimized on large image databases for tasks such as object classification. Instead, we optimized a naïve network starting from scratch to directly predict the recorded activity in the voxels of a given higher-order ROI, thereby bringing neuroscience data directly to bear on the model development process. Thus, the network learned to represent images in a way that is optimal for predicting voxel activity in that ROI, capturing only the relevant dimensions of variance and the tuning of the voxels. We trained a separate model for each of four ROIs: FFA, EBA, VWFA and RSC. We capitalized on the natural variation in the rich Natural Scenes Dataset (NSD) [32] to train these models directly on stimulus-response pairs from a wide range of naturalistic scenarios. Notably, the dataset contains complex and sometimes crowded images of various everyday objects in their natural contexts at varied viewpoints. The stimulus set is thus more typical of real-world vision, which permits the characterization of neural representations and computations in ethological conditions.

### Accurate predictions on complex, cluttered scenes; rapid generalization to new subjects

Our models learned deep convolutional feature spaces that are shared across thousands of voxels over multiple subjects. We utilized a rotation-equivariant Convolutional Neural Network (CNN) architecture to learn these feature spaces directly from fMRI data. This architectural design choice enables the model to learn identical features at multiple orientations and spatial locations, mimicking the response properties of neurons in early and intermediate visual areas, which are known to capture similar features like edges and curves but at different orientations and locations in the visual field [40, 41, 42, 43]. We employed a linear *readout* model on top of this feature space to predict the responses of individual voxels in the ROIs. The linear readout was *factorized* into *spatial* and *feature* dimensions following popular methods for neural system identification in mouse visual cortex [44]. This factorization separates receptive field location (i.e., what portion of the visual space is the voxel most sensitive to?) from feature tuning (i.e. what features of the visual input is the voxel sensitive to?). Sharing the entire representational network across subjects, sharing convolutional filters weights across visual field locations (translation equivariance) and orientations (rotation equivariance) and a factorized readout jointly enable the sample-efficient training of the response-optimized models (see Appendix D). Our baseline is a task-optimized model with a similar number of convolutional layers trained to perform object classification on the large-scale ImageNet dataset (see Fig.1 and the Methods section for details). To assess the prediction performance of this model, we used the standard methodology of modeling the voxel response as a linear weighting of the task-optimized network’s output units (from the best-performing layer that is determined using a separate validation set). Further, just as in our response-optimized models, this linear mapping is factorized into spatial and feature dimensions as this was found to significantly improve performance over the traditional non-sparse readout method.

**Figure 1:**
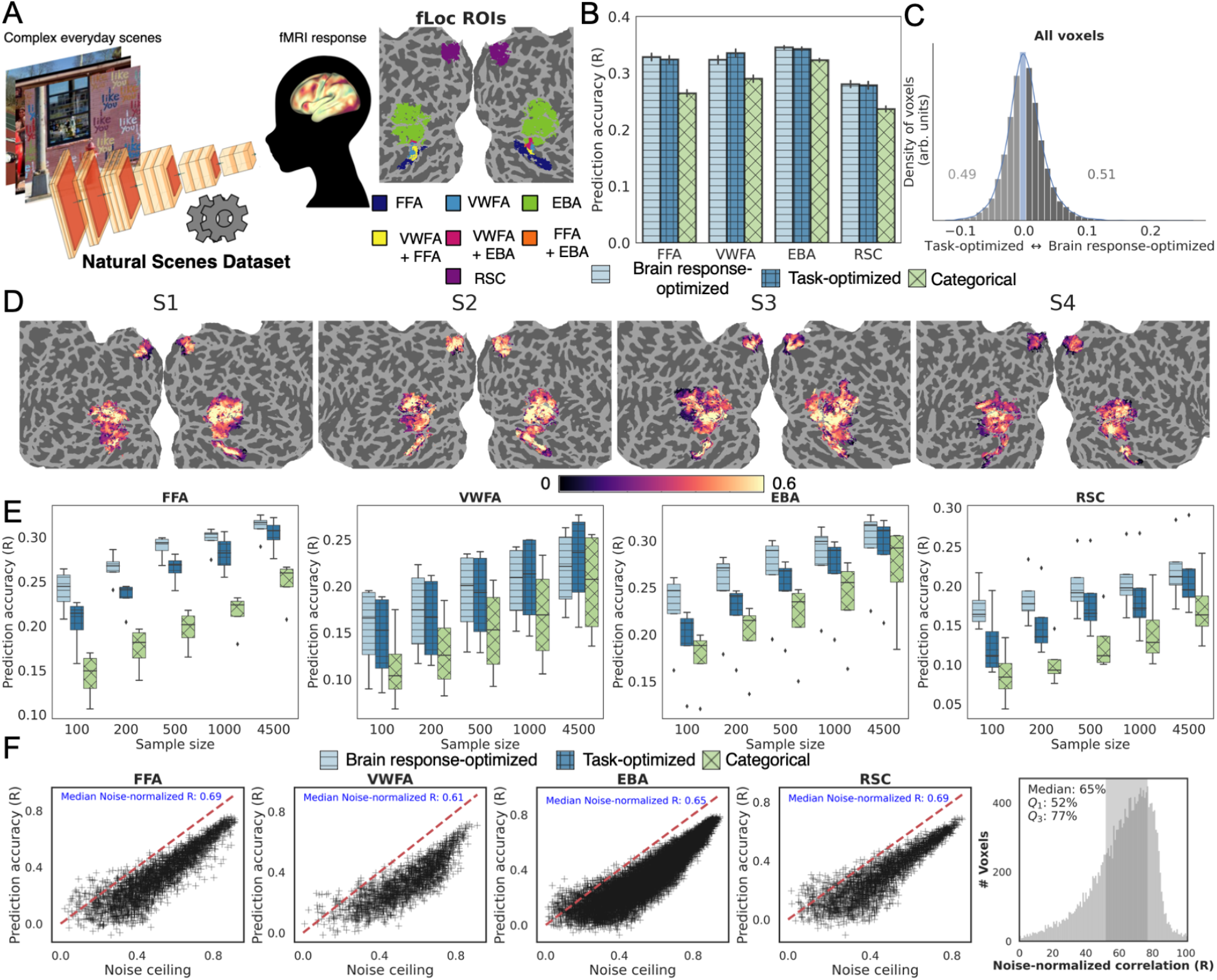
Performance of response-optimized and baseline models. **A** Schematic of the experimental paradigm and a cortical flatmap of the 4 visual ROIs studied here, which have some noticeable overlaps. **B** Prediction accuracy of the proposed and baseline models (a task-optimized model trained on object classification on ImageNet and a simple categorical model) as estimated by the Pearson correlation coefficient (R) between predicted and held-out responses for the same subjects on which the models were trained. Bars indicate a 95% confidence interval over all voxels of all subjects obtained via 1000 bootstrap samples. **C** Voxel-wise distribution of the difference between prediction accuracies of the response-optimized and task-optimized models. The inset shows the proportion of voxels that are better predicted by each. The response-optimized models achieve parity with the task-optimized model trained on a million ImageNet images (no difference was found through a two-sided permutation test, p*>*0.01 for all 4 ROIs). **D** Cortical flatmap illustrating the prediction accuracy achieved by the response-optimized model in all voxels of the four ROIs. High predictive accuracy (*>*0.6 unnormalized correlation) is achieved within large swathes of these ROIs. **E** Generalization performance of all models to new subjects as assessed by varying the amount of stimulus-response pairs used to train the linear readout. Response-optimized models generalize much more efficiently to novel subjects than task-optimized or categorical models. **F** Un-normalized prediction accuracy (R) of every voxel against the corresponding noise ceiling. Noise-normalized prediction accuracy is reported in the inset. Much of the variance in predictive accuracy across voxels is driven by their noise ceiling. Response-optimized models attained approximately 61-70% of the noise ceiling, functioning as one of the most quantitatively precise voxel-level models of these higher-order regions.

We compared the performance of response-optimized and task-optimized models (see Fig.1). The models are trained jointly on four subjects and their performance is estimated on a held-out set of 1,000 images seen by these same subjects. For these subjects, the response-optimized models attained approximately 65% of the noise ceiling (median correlation), with noise-normalized correlations lying between 52-77% for 50% of the voxels, yielding one of the most computationally precise voxel-level models of these higher-order regions. Further, response-optimized networks achieve parity with task-optimized networks on the same subjects (FFA: p = 0.274, VWFA: p=0.014, RSC: p=0.709, EBA: p = 0.510. p-values calculated by a two-sided permutation test with N=1,000). Consistent with results previously reported using recordings from non-human primates [21], our models are far more predictive than the category ideal observer model which employed the category membership of labeled objects in the image (another, simpler baseline than task-optimized models).

Next, we assessed how these predictive models generalize to the remaining set of four subjects that were not used to train the model. For this analysis, we train each network to predict activity for the remaining subjects by only optimizing the weights of the final linear readout while keeping the rest of the network fixed. We vary the amount of stimulus/responses pairs from the new subjects to train the readouts, from only 100 samples to a large set of 4,500 stimulus-response pairs. The influence of neural dataset size on predictive accuracy can complement prediction performance when evaluating the quality of two competing models. We consider the best representation to be the one that enables the most sample-efficient learning of the readout model for new subjects. Importantly, the difference in performance between response-optimized and task-optimized networks becomes even more striking as we limit the stimulus-response pairs (see Fig. 1.E). In FFA, EBA and RSC, the average performance for the response-optimized networks is already at more than 78% of its final value after just 100 training samples, compared with 60 − 67% and 50 − 65% for the task-optimized and categorical models respectively. In FFA and EBA, we need 500 samples for the task-optimized network to achieve a comparable performance to the response-optimized network with 200 samples. This remarkable generalization of response-optimized networks suggests that they are able to sufficiently constrain the space of possible solutions in the right manner so that the readouts for new subjects can be learned with few samples.

### Network dissection reveals selectivity, tolerance and clutter-invariance

Here we demonstrate that the high prediction accuracy and generalization of our hypothesis-neutral response-optimized networks do not come at the cost of model intelligibility. Instead, we show that the response-optimized models possess both empirical and aesthetic virtues, being computationally precise and elegant at the same time. Notably, features that emerge in the trained networks result from optimization to match ROI responses. After training the networks, we can probe their learned features, understand the computations they perform and, consequently, understand the characteristics of the ROI responses they model.

We adapted the recently proposed technique of network dissection [35, 36] to generate “verbal” explanations for the responses of different voxels. The technique measures the degree of alignment between a voxel’s response properties and an extensive visual concept dictionary, spanning objects and fine-grained visual concepts like parts of objects, colors, materials, and textures. We quantified the agreement between each concept and individual voxel using the Intersection over Union (IoU) metric following [35]. The IoU metric is computed on an independent, large-scale natural image database comprising a diverse set of real-world environments (indoor, urban, and natural). An alignment is computed between two maps for each image in the database. The first map indicates the high-level concept corresponding to each pixel in the image (pixel-level labeling is performed by humans or a high-performing segmentation network). The second map indicates the spatial regions within the image that are highly activated by the convolutional filter corresponding to a voxel (see Methods section for further details). After computing the alignment across images, the result is an alignment value for every voxel-concept pair. It is essential to note this methodological framework’s subtle yet profound implications. Previously, response profiles have been primarily defined using image-level category labels. Our approach enables us to model voxels as convolutional filters and systematically identify the image properties that they respond to without an a-priori hypothesis specification. Our approach characterizes not just ‘which’ images activate a particular brain voxel but also ‘what’ in those images drives the response, providing a rich characterization of neural responses to crowded natural scenes. For the top 20 concepts in each ROI, the median IoU is shown across all voxels in that ROI (Fig. 2B). Following [35], we used a stringent IoU threshold of 0.04 to detect ‘matching’, i.e., if a voxel’s corresponding filter exhibits a high agreement with a concept map (exceeding 0.04 IoU threshold), we classified that particular voxel as *detecting* or *encoding* that concept. Fig. 2C shows the concepts for which a high agreement is found and the count of the voxels that achieve a high agreement with that concept (each voxel is only counted once, against the concept with the top IoU). As a point of comparison, Fig. 2D shows the same measures for untrained networks having the same architecture as response-optimized models, but with random weights. Finally, Fig. 3 shows a visualization of the part of highly activating images that leads to the high activation. For each ROI and its preferred concept (according to the median IoU metric), the five top voxels were chosen, and for those, the top 100 images activating images were picked from the large scale dataset. For randomly selected images from this set, the area that leads to maximum activation of the voxel is shown in Fig. 3. 2

**Figure 2:**
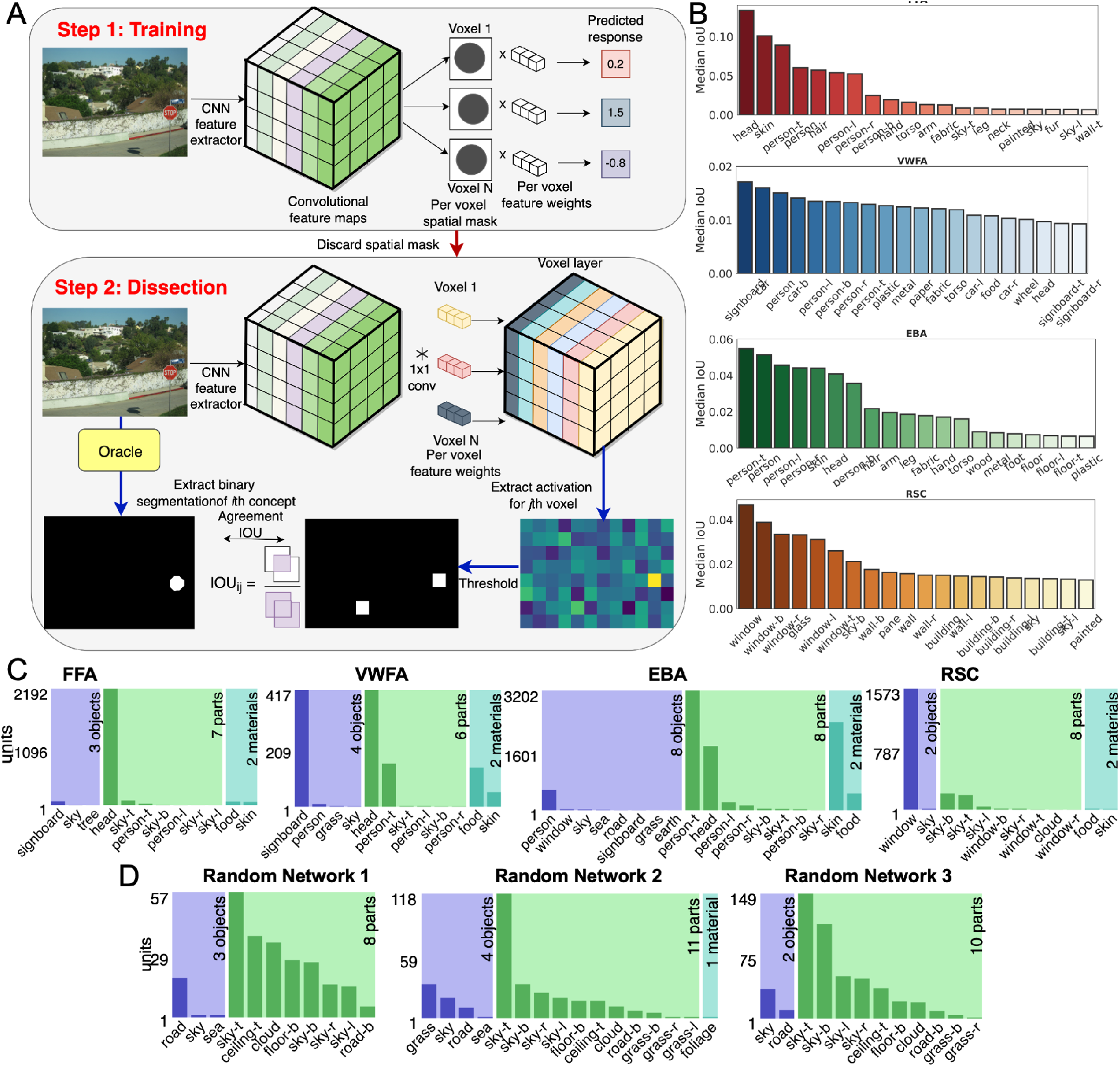
Selectivity revealed by network dissection. **A** Schematic of the model conversion procedure used to obtain the dissection results. The spatial mask was discarded and the learned feature tuning of every voxel was employed to create an additional 1×1 convolutional layer, so that every voxel is represented by an independent unit in this convolutional layer. **B** demonstrates the median IoU metric across all voxels belonging to an ROI for the top 20 visual concepts rank ordered by median IoU. The top concepts for FFA, VWFA and EBA discovered using this hypothesis-neutral approach (‘heads’, ‘signboards’ and ‘person’ respectively) align remarkably well with the widely held selectivity of voxels in these regions. **C** shows the matched concepts for every response-optimized model, i.e., the number of units in the ‘voxel’ layer that showed high alignment (stringent threshold of IoU*>*0.04) with a human-interpretable visual concept. Multiple units (i.e., multiple voxels) are associated with the same high-level concept. Contrary to object recognition networks which are trained with explicit label supervision and which show a broad diversity of detectors [35, 36], the matched visual concepts in these response-optimized networks are highly specific and aligned with the previously hypothesized functional role of these ROIs. **D** shows the matched concepts for 3 networks with the same architecture as response-optimized models but random weights. The detection of these concepts (e.g. sky, grass, road) is likely driven by low-level cues, like color, instead of complex features.

**Figure 3:**
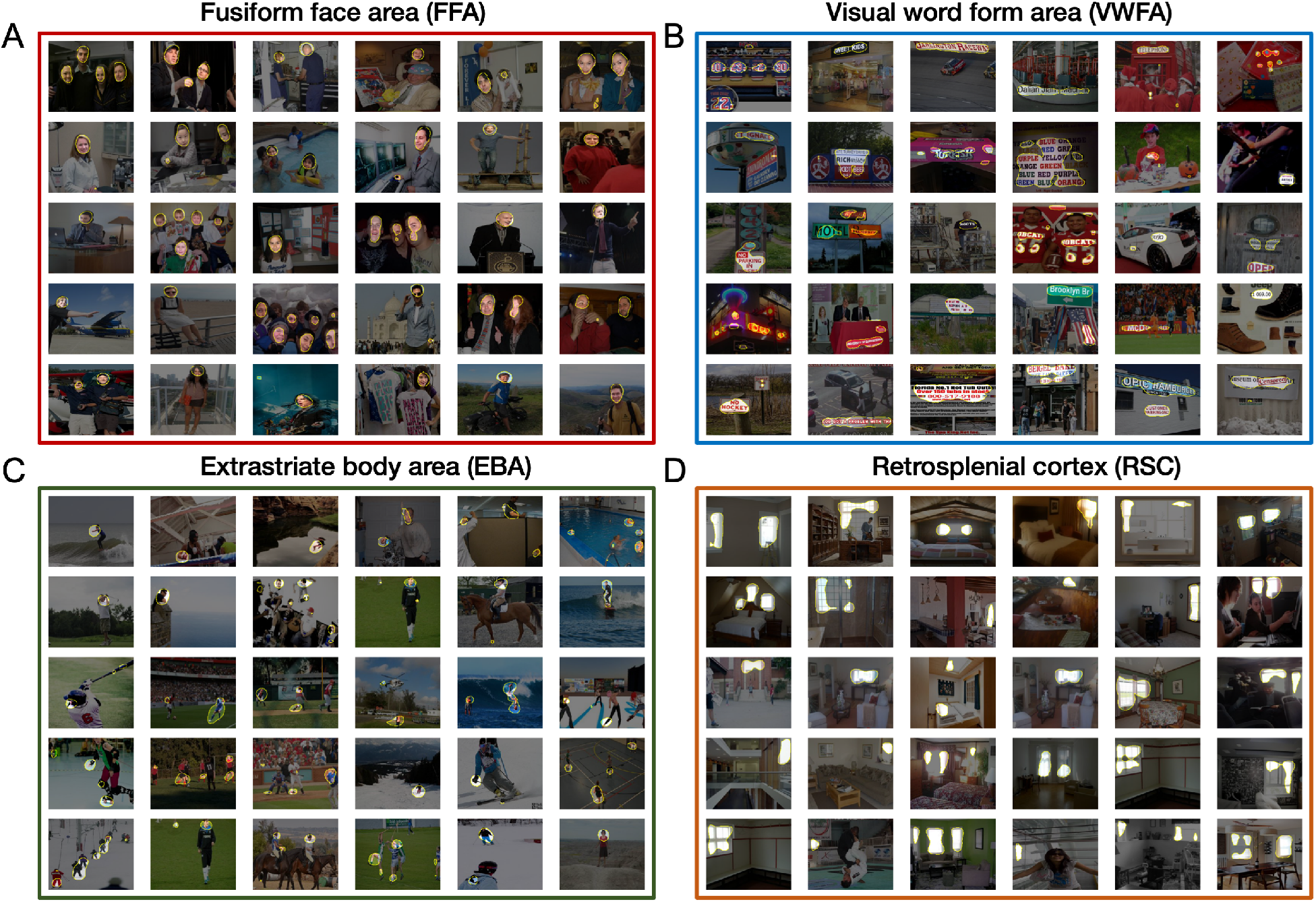
Segmentation maps resulting from network dissection. Activation of a single voxel (converted to a 1×1 convolutional filter) in response to an input image is visualized as the region in the input space that elicits the highest (top 1% quantile level) activation in the corresponding filter output. Top five voxels as ranked by the IoU with the preferred concept for every ROI (‘heads’, ‘signboards’, ‘person’ and ‘windows’ for FFA, VWFA, EBA and RSC respectively) were identified and input images were randomly selected for each voxel among the top 100 most activating images for that particular voxel to maximize diversity across voxels. Each row corresponds to a distinct voxel within the respective ROIs.

Fig. shows that the FFA network’s favorite concept is ‘head’ (the large scale dataset didn’t have a face label, and faces are labeled as parts of heads). The head concept has the highest median IoU of 0.125, more than other top concepts for FFA and the other ROIs. Some FFA voxels predictors have IoU as high as 0.20 with the ‘head’ concept. These predictors can be contrasted with the ‘head’ detectors (model units with IoU*>*0.04 against the ‘head’ concept) that emerge spontaneously in the last convolutional layer of a standard AlexNet trained on image categorization (ImageNet)^1^. In this task-optimized model, the best detector yields an IoU of 0.15 against the ‘head’ concept maps with the median IoU among all head detectors being 0.08. Thus, the face selectivity of units in the FFA model even exceeds the face selectivity of units in models trained explicitly with semantic labels on a million images. Fig. 3[A] illustrates how the chosen voxels in FFA act as head (face) detector, with the parts of images driving the predicted response being almost exclusively faces.

The VWFA network has high median IoU with signboards (containing letters) even though the number of images with writing is small in the NSD dataset. Fig. 3B shows that signboards with very different lettering and signs (different backgrounds, fonts, styles, scales, colors, orientations etc) drive the predicted response in the top voxels, even though the scenes are often cluttered with a myriad objects simultaneously. This result replicates the highly invariant orthographic processing in the VWFA [45]. The VWFA also has high alignment with the concepts of head and person, which is due to the anatomical overlap of localized VWFA with FFA, discussed in a following section. EBA has high IoU with people, head and skin. The IoU with the people concept is highest, highlighting that EBA is tuned for body parts. Interestingly, for RSC, we observe a large number of ‘window’ detectors after applying the dissection procedure. Navigational affordances are important for scene perception, and windows might be particularly indicative of such affordances in indoor scenes and critical to functional scene understanding (for e.g., windows indicate an obstructed path where movements are blocked) [46]. Other concepts evoking high response within the RSC (e.g., ‘wall’, ‘glass’, ‘building’) relate to scene perception and navigation, in contrast to concepts discovered within the models optimized for the other three ROIs. These results highlight the differences in selectivity between the visual areas. Despite no access to category labels during training, our networks gained a strong semantic selectivity for high-level visual concepts and this selectivity remained tolerant to changes in category-orthogonal nuisance variables (e.g., size, viewpoint, clutter etc.), as shown in Figure 3.

### Maximally exciting images reveal structured high-level features consistent with previously hypothesized ROI role

In classical neuroscience, identifying the optimal visual stimuli (the peak of the tuning curve) for different neurons has been instrumental in understanding neuronal selectivity and its contribution to perception. Our models permit following this approach for higher-order visual areas where the optimal stimulus can be arbitrarily complex. We performed an unconstrained optimization over input noise to discover input images that result in maximal (predicted) excitation of individual voxels. We refer to these images as the maximally exciting inputs.

By performing unconstrained optimization, we are not restricting the space of maximally activating inputs to the naturalistic domain. While optimization without naturalistic constraints may impose its own set of challenges including generation of hard-to-recognize visual features, visualization with regularization or naturalistic constraints may not be truly faithful to the model. We favored the former approach and did not restrict the space of maximally exciting inputs to the naturalistic domain. Instead, we let the optimization process evolve complex visual inputs without constraints.

Fig. 4 shows that maximally exciting images for different voxels in the same ROI capture very similar visual properties. Out of all the features that could emerge from unconstrained optimization in a network trained on cluttered natural scenes, from simple features such as rounded shapes or eyes to possibly more complex high-level features, face-like images with overlapping small and large circles almost exclusively pop up for all FFA voxels, providing a strong support for the hypothesis that full ‘face’ features lead to increased activation in FFA. Similary, for EBA, we observe elongated curved shapes, loosely similar in form to body parts such as arms or legs. Maximally exciting inputs for the VWFA resemble orthographic units comprising curves and lines of different stroke widths like the visual form of letters. Finally, the maximally exciting inputs for voxels within RSC are reminiscent of windows (in accordance with the Network dissection results) in different reference frames, which may be linked to RSC’s role in spatial cognition [47]. We also observe rectilinear features in the maximally activating images for RSC, consistent with the previously found rectilinear preference of scene-selective areas [48], while the EBA and the FFA only captured curvilinear features. Therefore, our results provide further evidence for this hypothesis by learning the model that best predicts the data instead of starting *apriori* with the hypothesis.

**Figure 4:**
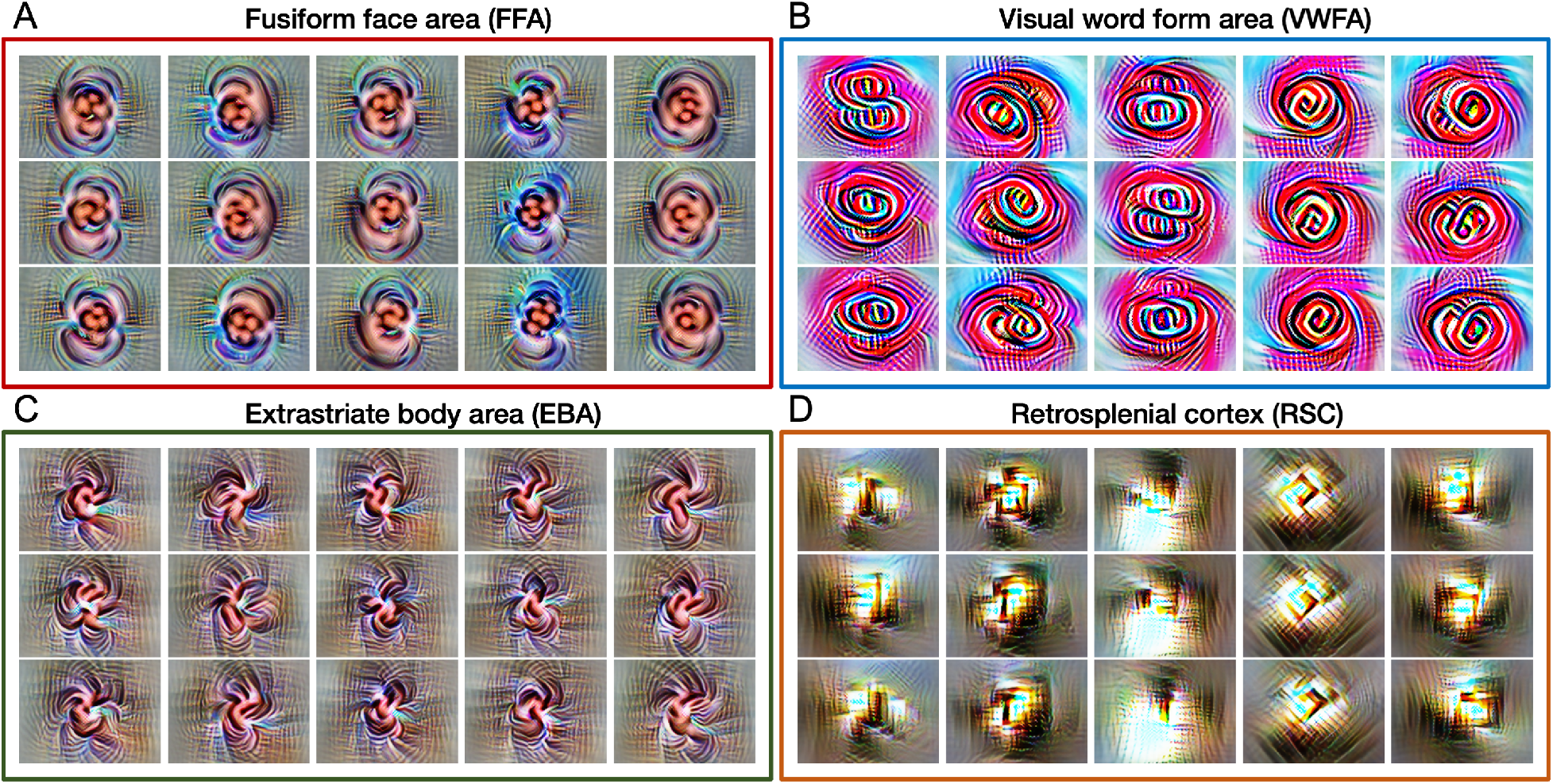
Maximally exciting inputs obtained through optimization. Maximally activating images as discovered by the optimal stimulus identification procedure, wherein an unconstrained optimization is performed over random input noise to discover the input that would result in maximal excitation of individual voxels. Images are shown for 5 randomly selected voxels per ROI starting from 3 random initial points per voxel. Each column is a randomly chosen voxel and each row is a different initialization. This structural analysis reveals that the features that drive individual voxels (the *visualizations*) are consistent with the *verbalizations* for those voxels as expressed by the network dissection procedure. More specifically, face-like visual forms, curved features akin to orthographic symbols, skin-colored shapes reminiscent of some body parts and window-like rectilinear features emerge spontaneously for FFA, VWFA, EBA and RSC respectively.

### End-to-end models capture tuning differences between voxels in the same ROI

To investigate if the proposed models indeed capture meaningful differences between voxels, we computed *spatial generalizability matrices* by correlating the predicted response of each voxel against the measured response of every other voxel to obtain an N×N correlation matrix for shared models, where N is the total number of voxels across all participants [49, 50, 51]. These matrices reveal the similarity of the tuning of each pair of voxels. To account for higher variability in measured versus predicted response, we normalize the rows and columns of this correlation matrix following [52]. The diagonal dominance in these identifiability matrices (Fig. 5A) suggests that predicted responses are most similar to the same voxel’s measured responses, which indicates that all models are successfully able to capture meaningful voxel-level idiosyncracies to some extent. The generalization matrices reveal the presence of several distinct clusters (at least two) for all ROIs, such that the models of voxels in one cluster are highly predictive of responses of voxels in the same cluster (both within, and across participants), but do not generalize to other clusters. Importantly, this clustering structure is prevalent across participants (specifically for FFA, EBA and VWFA, with more variability for RSC), indicating a shared organization. We leave a thorough characterization of the differences between these clusters for future work.

**Figure 5:**
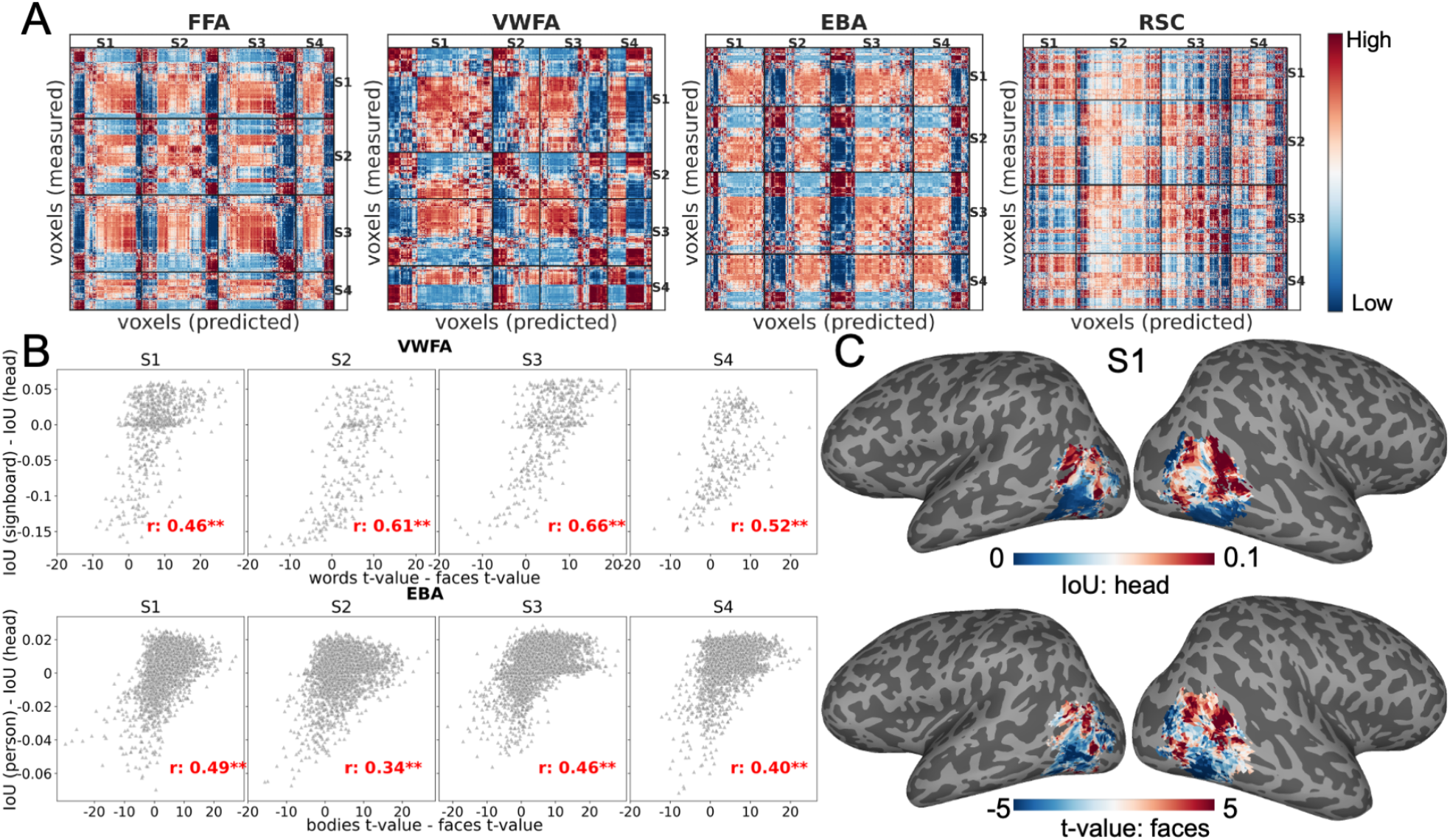
Voxel-level tuning characterization. **A** shows *spatial generalization matrices* for every ROI computed by correlating the predicted response of every voxel against the measured response of every other voxel belonging to the same ROI (within and across subjects). Black lines mark subject boundaries. **B** Scatter plot depicting the difference between IoU of the preferred category (loosely, ‘signboard’ and ‘person’ for VWFA and EBA, respectively) and faces against the difference between the corresponding t-statistic values estimated by the functional localizer experiment. Each point is an individual voxel belonging to the same ROI. The inset indicates the correlation coefficient and significance was estimated through a two-sided test. **C** Cortical surface plot of face selectivity as measured by the localizer experiment against the IoU with the ‘head’ concept as quantified by the dissection procedure for one subject (Similar plots for remaining subjects are shown in Supplementary Fig. S1).

Next, we examined voxels that were strongly selective for a different semantic concept than the conjectured preferred category for their visual area. We specifically examined the ‘head’ selective model neurons identified by the network dissection procedure within VWFA and EBA. We assessed the correlation between the quantitative agreement of these voxels with the ‘head’ concept over the ‘signboard’ or ‘person’ concept (IoU, as evaluated by our dissection procedure) and the degree of their face-selectivity over selectivity for words or body parts (as quantified with the independent functional localizer experiment). Our results suggest a very strong correspondence between the two (Pearson’s R∼0.34-0.66, p*<*0.001 for all 4 subjects and both comparisons, Fig. 5B), despite the very different experimental paradigms involved in the two quantifications. Further, in addition to comparing this relative selectivity, we also compared the absolute ‘head’ selectivity of voxels in EBA and VWFA against the face-selectivity (t-value) measured with the localizer experiment. Again, we see a striking pattern of similarity in these estimated and measured values (Pearson’s R∼0.6-0.7 (p*<*0.001); (see Fig. 5 and Supplementary Figure S1). Thus, our computational models successfully capture the spatially overlapping representations of semantic categories and graded functional organization within human extrastriate cortex.

### High selectivity persists in ‘face-deprived’ and ‘body-deprived’ networks

The strong semantic selectivity in our response-optimized models leads to an important question: are the ROIs they model simply functioning as detectors for their preferred category? In this case, then training for example a neural network to predict FFA activity would be equivalent to training this network with images associated with a label that indicates the presence of a face. An alternative hypothesis is that category-selective ROIs are sensitive to visual properties that are typical of their preferred category, even in the absence of that category. In this case, training a neural network to predict FFA would provide it with a more complex signal that allows it to learn the sensitivity of the FFA to those visual properties.

To differentiate between these alternatives, we focused on FFA and EBA and trained response-optimized models with the same architecture above but with a visual training diet entirely deprived of images containing the ‘person’ category. Surprisingly, we find that, despite not seeing *any* image with human faces or bodies during training, the networks optimized to predict FFA and EBA retain their categorical selectivity (Fig. 6). Supplementary Fig. S6 further shows that these results hold even when further removing any images with animals from the training set. The training diet deprivation did decrease the agreement of model voxels with their respective preferred category (e.g., in FFA, the median IoU with ‘head’ dropped from ∼0.13 in the non-deprived network to ∼0.08 in the deprived network), indicating a slightly reduced selectivity (Fig. 6). However, the preferred concept was still overwhelmingly ‘head’, and the resulting segmentation pictures look qualitatively similar to those in Fig. 3. In other words, we found that networks trained with a visual diet deprived of their preferred category could still extrapolate the responses to the preferred category.

**Figure 6:**
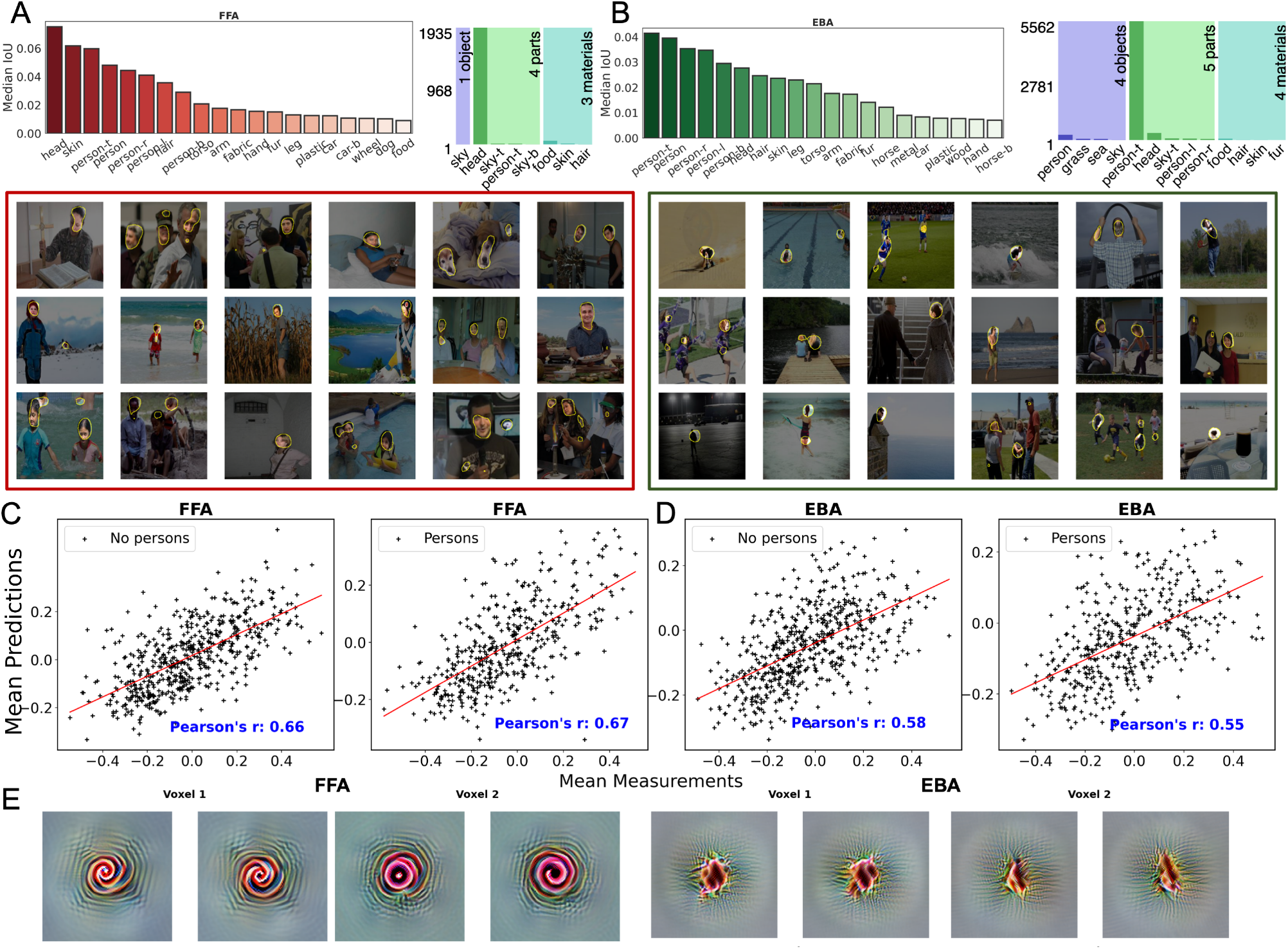
Selective deprivation in visual diet. **A** and **B** show results of the dissection procedure applied to response-optimized models of FFA and EBA respectively, when trained using a visual diet entirely deprived of the ‘person’ category; Left panels show the median IoU computed across all voxels in that ROI against the top 20 concepts identified using the median IoU metric. Right panels show the matched visual concepts for the respective ROIs. An IoU threshold of 0.04 is applied to detect matching. Activation of 3 randomly selected concept detectors in each ROI model to top images are shown below. **C** and **D** depict the mean measured response across all voxels for every test image against the corresponding mean predicted response, separated into ‘persons’ and ‘no persons’ categories for FFA and EBA respectively. Pearson’s correlation coefficient between the mean predicted and measured values is reported inside each scatter plot. Similar results for models trained with a visual diet deprived of all animate categories are shown in Supplementary Figure S6 and voxelwise results are shown in Supplementary Figure S9. **E** depicts the most activating images, as discovered by the optimal stimulus identification procedure, for two randomly selected model neurons emulating brain voxels (from the deprived response-optimized models) using 2 random starting points. Concentric circles emerge for FFA voxels, consistent with previous studies demonstrating strong FFA activation to concentric gratings [53]; skin colored patterns emerge for the EBA voxels.

Further, we observe in Fig. 6C and D that models do not incur a severe loss in prediction performance when data from a new domain (i.e. ‘faces’ and ‘bodies’ category) is presented at test time. Actually, the FFA model achieves even slightly better prediction performance on held-out images with faces. This example of systematic generalization in the developed response-optimized models is also interesting from the perspective of modern deep learning, which is often criticized for its failure to generalize in this systematic, out-of-distribution way. The ability of these models to generalize to new domains further validates their proposed usage as virtual stand-ins for fMRI experiments, supporting their ability to act as *model organisms* for large-scale fMRI experiments and revealing selectivity for stimuli beyond those encountered in the training set.

### Models reveal key functional distinctions between different regions

Our results indicate that response-optimized models capture complex tuning properties relevant to their preferred category but not uniquely exhibited by it. Next, we hypothesized that our models *meaningfully* discriminate between stimuli belonging to their preferred set. More specifically, we tested existing accounts of functional specialization which implicate FFA in face perception, particularly face identity discrimination [54, 55], RSC in spatial cognition [47], EBA in perception of the shape and posture of bodies [56], and VWFA in visual perception of symbols, like letters and digits [57].

We first performed a face discrimination task using the response-optimized models of all four ROIs. For each model, we extracted the predicted voxel-wise response for a small subset of facial images from the CelebA dataset, comprising 20 identities, each with 100 train, 30 validation and 30 test images [58, 59]. We compared the face recognition accuracy of these model predictions against the recognition accuracy of a general-purpose representation from a CNN trained to perform image classification on the large-scale ImageNet dataset. We also compared the face recognition accuracy of these models with that of a representation from a network trained explicitly to do facial recognition on a large set of VGG face identities [60]. For each of the representations above (the four ROIs’ predictions and the general-purpose and face specialized representations), we trained a linear function to predict the CelebA identities. The resulting FFA predictions significantly outperform the predictions from all other ROI models (discrimination score of 85% compared to 79-80%), even outperforming the highly transferable representation of ImageNet trained networks [61] at 78%. We note this performance still falling short against features from networks trained to discriminate a large number of identities at 96%. This result provides additional evidence for the role of FFA in face identification and shows the ability of our model to pick up these functional capacities.

Next, we performed a room layout prediction task using the voxel-wise predictions generated by the four ROI models and the representations generated by the general-purpose CNN trained on ImageNet. The objective of this task is to predict the correct layout type from the 11 categories described in [62] defined using a keypoint-based parameterization (Figure 7[B]). The dataset comprises 4,000 natural scenes across diverse indoor scene categories from the SUN database [63], that are split into sets of 3200 training, 400 validation and 400 test images. We followed the same procedure for quantifying the layout estimation accuracy as before, i.e., training a linear classifier on top of each representation. Here, the results follow a different trend (Figure 7D). The RSC predictions outperform the other ROI models. They perform just as well as the ImageNet trained representations in solving this task, even though the RSC network was trained with ∼35,000 images and their associated brain responses in contrast to the ImageNet network training size of a million images. This result strongly supports the role of RSC in scene understanding, particularly when relevant to spatial navigation, and again highlights our model’s ability to transfer to these tasks.

**Figure 7:**
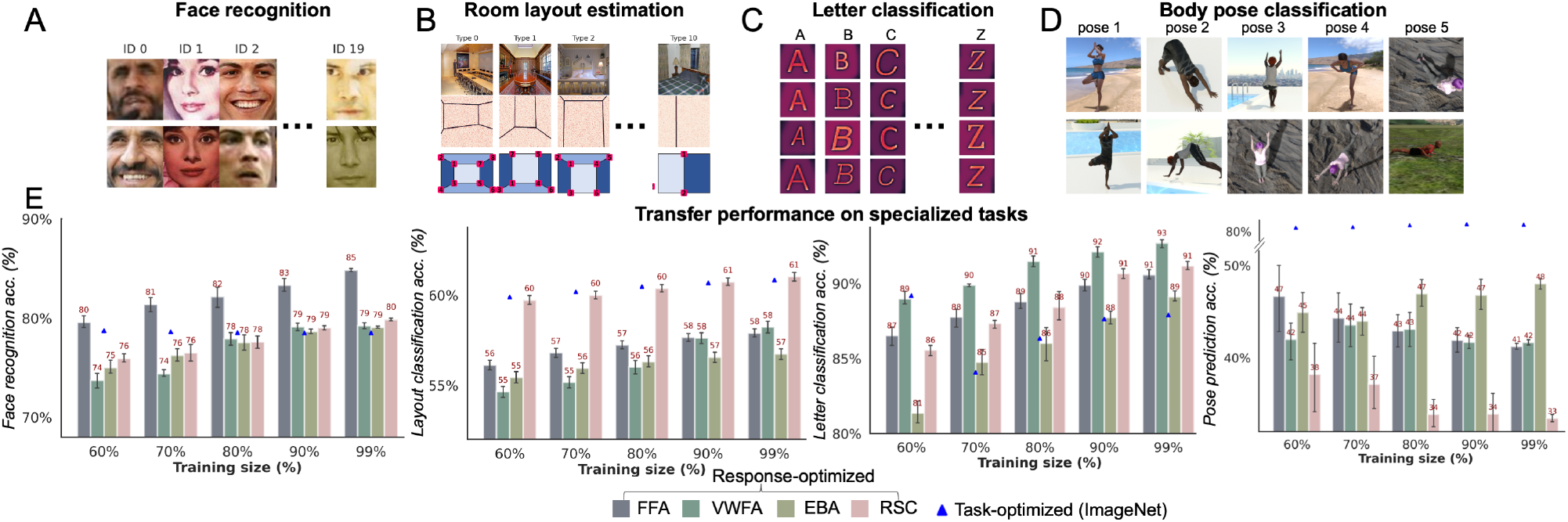
Task performance of neural representations. **A**,**B, C** and **D** show sample stimuli along with their classification labels for the four visual discrimination tasks considered in this study, namely face identity recognition, room layout, letter and body pose classification. In **B**, the ground truth layout planar segmentation of indoor scenes in the room layout estimation task are depicted below the respective scenes and the definition of their corresponding room layout types are illustrated in the bottom panel. **E** depicts the transfer performance of neural representations from response-optimized networks of each individual ROI on the four tasks. Bars indicate the standard deviation across five cross-validation folds. Intriguingly, representations from FFA achieve the best face individuation accuracy, RSC representations perform best on the layout estimation task and those from VWFA and EBA perform best on letter classification and body pose classification respectively, remarkably consistent with (and providing strong evidence for) the previously hypothesized functional roles of these visual areas.

Next, we followed the same procedure as above and trained a linear classifier on top of different model representations to perform a letter classification task. Here, the goal is to predict letter identities (A-Z) from images of printed letters created using different fonts, styles, sizes and position effects [64] (examples shown in Fig. 7C). The VWFA response-optimized model consistently outperforms the remaining response-optimized and task-optimized models (Figure 7D), providing computational support for the ability of VWFA responses to enable letter perception. We note that differences between models are small, possibly because of the substantial overlap between ROIs due to the liberal thresholds (t*>*0) applied on functional localizer data. Following a similar procedure, we quantified the ability of different model representations to support body pose classification on a computer graphics (CG)-generated body pose dataset containing exemplars from 5 well-known yoga poses [65].

On this task, we find that the EBA model consistently performs on par with or exceeds the performance of the remaining ROI models. Here, however, we observe a significant gap between the performance of the best response-optimized model and the ImageNet-trained task-optimized model, possibly due to the richer variation of different body postures in the ImageNet dataset as compared to NSD stimuli, which may allow efficient transfer to this task. All response-optimized models, however, perform significantly above chance (20%) on this challenging dataset.

We note that while each of these regions has been stipulated to fulfil a specialized function before, thus far, no study has shown evidence of this task specialization from the pattern of neural responses in these regions directly (see [5] for an exception) or computational models of these regions. In fact, evidence from task-optimized models suggests the contrary. Deep neural networks trained on face individuation perform worse than or on par with networks trained on object categorization in capturing face representations in the macaque [66] and human brain [67], posing a challenge to domain-specific functional accounts of face-selective regions. How do we reconcile our results with these findings? In modern DNNs, task-optimization and training datasets are often intertwined due to the limited availability of datasets that contain multiple types of learning signals (i.e. labels). This coupling confounds interpretations and models trained on ImageNet often surpass other models due to the rich coverage of the naturalistic stimulus space in this dataset over others, which can compensate for lack of sufficiently powerful priors; these ImageNet-trained object categorization models continue to yield best response predictions for all ventral visual regions alike [24]. In contrast, the response-optimization approach offers an intriguing opportunity to probe functional distinctions among ROIs since these models can be trained on an identical image set and only the training signal (i.e., the brain response) can be varied between these models. Any differences in emergent function within the trained models can thus be more confidently attributed to the differences in response patterns between ROIs. Further, we note that category selective responses and functional specialization (fine-grained discrimination between exemplars of the preferred category) do not necessarily follow from each other. Selectivity is contingent on the extraction of features shared by all exemplars from the preferred category while fine-grained discrimination relies on features that distinguish each exemplar (each facial identity or each room configuration) from others. While selectivity is measured with respect to responses of single neurons or voxels, fine-grained discriminability is a measure of information contained within the population response. Selectivity and functional significance analyses thus do not equate from a computational perspective and offer a complementary purchase on our understanding of these regions.

### Hypothesis-agnosticity reveals a limited set of categories for which selective responses are observed in the ventral visual cortex

We note that our approach, thus far, was hypothesis-neutral in the sense of model construction; i.e., the models were unconstrained by any particular hypothesis, rather they were largely informed by the data. However, the selection of regions was nonetheless biased by apriori knowledge and their localization was guided by independent functional data measuring the brain responses to different categories of stimuli. We presented an approach to determine the selectivity of these regions and characterize their function in an impartial computational manner. The emergence of category-selective units in response-optimized models unbidden provided emphatic evidence for category-selectivity of these regions and suggested that these regions not only respond to low-level features most correlated with the preferred category but to the high-level category per se. Here, we take a step further and ask: can we discover category selectivities with an impartial approach that does not require ROI selection or localization? To answer this question, we performed a retrospective analysis: we trained a response-optimized model on responses from the entire ventral visual cortex (28,910 voxels) across all subjects. The resulting model is able capture the structure of ventral visual stream responses, as evidenced by a reasonably high response prediction accuracy (median noise-normalized R ∼ 59%). We then applied the network dissection procedure on units within the optimized model. The results, shown in Figure 8, highlight the categories previously believed to elicit selective responses, namely heads (or, faces), signboards (or text) and skin, which reflects selectivity for the ‘bodies’ category. A small number of units appear selective for elements such as sky and window, which can be attributed to processing scenes. Moreover, we also observe single-unit selectivities for the ‘food’ category, for which neural selectivity was also reported in recent works [37, 38, 39], validating our data-driven approach. The non-existence of units aligned with any of the thousands of other semantic concepts beyond these categories in the dissection concept dictionary (e.g. car, door, chair, plastic etc.) also serves as evidence *against* the existence of selectivity for those categories of stimuli in the ventral visual cortex.

**Figure 8:**
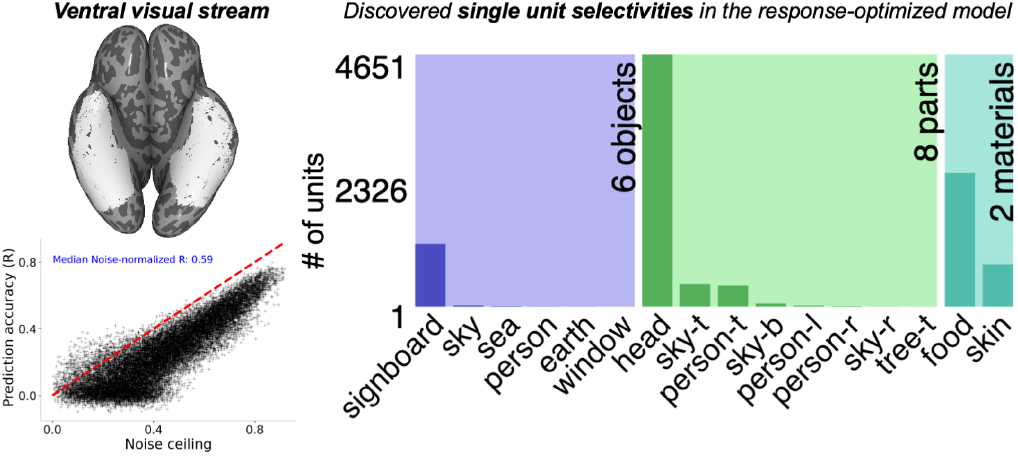
Dissection (right) of a response-optimized model trained to predict responses across the entire ventral visual stream (left, shown for one participant). Bottom left plot shows the un-normalized prediction accuracy (R) of every ventral visual stream voxel against the corresponding noise ceiling. Noise-normalized prediction accuracy is reported in the inset.

## Discussion

Here, we exploited data-driven, hypothesis-neutral deep neural networks to model the responses of higher-order visual ROIs, specifically FFA, EBA, VWFA, and RSC. Through a large, rich stimulus set afforded by the NSD, we offered new evidence that generalizes decades worth of hypothesis-driven results to ethologically valid empirical settings, and further refines these hypotheses.

### Brain response predictivity

We found that response-optimized deep neural network models—trained solely with supervision from fMRI activity—accurately predict activity related to new images in multiple visual category-selective ROIs. The performance of these models rivals the predictive performance of state-of-the-art task-optimized models used to predict brain activity [21]. Many model networks can be consistent with brain activity data as demonstrated by the number of recent papers in this area [68]. It can be argued that a good representation would allow for efficient learning (in terms of the number of data points required) of a linear predictor of brain responses. Thus we ran an analysis in which we evaluated the trained models on the ability of their representations to predict data for new subjects while restricting the training set size. We found that the response-optimized models were better able to generalize to novel subjects at small sample sizes than task-optimized models. This result suggests that end-to-end optimization solely driven by response measurements can yield better correspondence to brain data than networks optimized on behaviorally relevant tasks with millions of images.

### Verbalization of voxel selectivity with network dissection

The current trend in visual and computational neuroscience uses naturalistic stimuli not amenable to parametrization and neural networks to model brain responses. This approach requires concomitant methodological advances to derive trustable conceptual understanding from brain response models. This view assumes that accurate and generalizable computational models of brain function can serve as a reasonable proxy to biological visual systems. Probing and interrogating these models may provide novel insights into the properties of their associated biological system. Here, we developed a systematic methodological framework to understand the tuning properties learned by our response-optimized networks. To reveal unidentified neuronal tuning properties, our framework relies on the recent progress in understanding neural networks using large-scale annotated datasets and network dissection procedures [35]. This alignment procedure capitalizes on the use of a factorized readout that disentangles the spatial (“where”) and feature (“what”) dimensions of the voxel’s response properties and fully characterizes the “what” dimension using dissection procedures applied on the ‘model neuron’ corresponding to the voxel.

Using complex stimuli like cluttered natural scenes containing multiple objects in context to probe neuronal response properties poses multiple challenges. If an image set that highly activates a voxel is identified, several competing interpretations can be ascribed to that voxel’s tuning function. There can be multiple low-level (e.g., color or shape) or highlevel (e.g., semantic) features consistent with the identified images. When stimuli become more complex, recognizing the consistent pattern implicit among any given image set becomes more challenging as visual features become increasingly tangled (textures, shapes, edges, etc.). The dissection procedure in this study helps simplify this problem by not only recognizing ‘which’ natural images elicit higher responses, but also systematically and quantitatively characterizing ‘what’ part of each image drives increased responses. With this approach, we can effectively verbalize the functional role of voxels in each candidate region. We demonstrated that emergent concepts in response-optimized networks are highly specific and aligned with each region’s previously hypothesized functional role. For example, we reported an exclusive presence of face detectors in the model for FFA. If the learned category-selectivity were due to low- or mid-level features correlated with the category of interest, those features should have become apparent. The dataset used in the network dissection analysis is one of the largest densely-annotated natural image datasets in computer vision research. If the selectivity of FFA to faces could be explained by the unique structure of eyes or skin-colored textures, response-optimized models would have highlighted these simpler features. The segmentation-based dissection procedure would have revealed this pattern of selectivity as it scores units against category-level labels and thousands of other concepts, including different textures and object parts (including eyes, nose, etc.). With our dissection procedure, we see an almost exclusive selectivity for full faces in the FFA voxels.

Our results also indicate that computational models can account for perceptual invariance, even though they are only optimized for brain response prediction. The models respond selectively to their preferred category despite tremendous variation in the precise physical characteristics of the preferred objects.

### Visualization of voxel selectivity with image synthesis

Next, we used an optimization-based image synthesis technique to construct the stimulus that causes synthetic neurons modeling individual voxels to activate maximally. Several recent studies have attempted to describe the tuning properties captured by DNN models of visual cortex with similar image synthesis algorithms [69, 24, 70]. These reports found such methods could control neural firing activity in mouse primary visual cortex and macaque V4 [71, 72]. However, several distinctions between foundational work in this direction and our results are worth consideration. Unlike some studies that use the hypothesis space of generative adversarial networks, we do not impose a ‘naturalness’ prior on the optimized images. We neither employ a task-optimized model trained on large-scale object databases to derive the features that map onto brain activity. Both these choices may bias the features to contain high-level semantic content. Since task-optimized models are trained with explicit category-level pressures, this bias can arise even if the neurons/voxels primarily encode low-level or mid-level features that are not naturalistic or neatly verbalizable. Moreover, existing fMRI studies apply such techniques at the ROI level [24, 69]. Even though our optimal inputs were synthesized de novo from random pixel noise to activate single voxels in our study, we find that the optimized images still contain human recognizable complex patterns consistent with the hypothesized selectivity of these voxels. The images appear to capture critical functional properties of voxels in these high-level visual regions. We argue that encoding models fitted directly to neural data with no prior training, offer a useful, complementary perspective for understanding the visual system as opposed to hypotheses-driven, task-optimized models.

Any set of features that emerge in empirically-estimated response-optimized networks are optimized to explain representations in the brain, and are not tangled with confounds from top-down constraints unrelated to neural activity.

### Specific versus non-specific mechanisms in shaping categorical selectivity

Next, we analyzed the role of visual experience in shaping response selectivity by training response optimized models with a visual diet completely deprived of faces or bodies. Despite this selective deprivation, units in models of FFA and EBA retained strong selectivity for their preferred semantic content. We posit that the models effectively became face and body selectors, which suggests that they could generalize their predictions to stimuli that were not included in training and learn which stimuli will induce maximal firing. The models could infer the preferred category of their corresponding ROI through training with a dataset devoid of this preferred category. The models could enable this generalization because of the properties of the representations of FFA and EBA voxels. These voxels appear to respond to visual configurations characteristic to faces or body parts, respectively. These configurations could be present in parts of images that do not correspond to a person but happen to have face-like (e.g., circles) or body-like (e.g., elongated, curved shapes) features. One potential mechanistic explanation is that attributes of these specific classes of visual stimuli (like faces and bodies) do not vary independently with the rest of the visual world and visual category-selective ROIs act like filters that constantly look for matches to their visual properties. While faces or bodies can activate these filters maximally, other visual patterns that resemble these categories can also activate the filters. For the FFA, these other visual patterns could act like pareidolia, or perhaps subtle pareidolia that is hard to detect by humans but that still activates the FFA.

The systematic generalization and extrapolation ability in our computational models suggests they can, in principle, discover functional specializations in underexplored parts of the visual system. Most claims regarding functional specialization and domain-specificity in the brain are grounded in hypothesis-driven contrast-based experiments, which might not include the best-fit stimuli for these regions. Even large datasets like NSD might exclude certain categories that contain key features for a given brain region. After training our models on under-explored regions, their generalization ability can aide in characterizing selectivity post-hoc.

Our results can also inform studies on statistical learning in artificial and natural systems. In fact, out-of-distribution generalization remains a key issue in machine learning. However, our model could robustly generalize to unseen categories. This ability reflects the strong semantic selectivity of the ROIs and their responsiveness to features characteristic of their preferred category even in the absence of that category. Another key question focuses on whether developing selectivity for faces requires experience with the unique structure of faces. Existing behavioral evidence from some face deprivation studies [73] suggests that face-processing abilities can persist without any face-specific experience. While subsequent studies have provided counter-evidence, this critical dispute still has no consensus. Our experiments highlight that face and body selectivity can emerge spontaneously in computational models with no face and body experience, and this selectivity is maintained across diverse naturalistic variations (see segmentation maps in Fig. 6). We do note the caveat, however, that our models were trained using supervision from brain regions that have experienced faces and bodies.

### Link between emergent representations and their functional roles

Goal-driven deep learning models of the visual cortex have paved a new way forward for understanding the representations and computations carried out by the visual system. They not just predict neural activity to novel stimuli, but also link it to complex visual behaviors in naturalistic domains due to their functional constraints. In fact, it is striking that biologically plausible representations emerge spontaneously in neural networks trained only with a handful of computational principles (gradient descent, convolutions etc.). While response-optimized models do not provide such a direct link to perception and behavior, here we asked if it may be possible to read out ‘behavior’ ensuingly from these models that are constrained solely with neural data, thereby investigating the functional role of different visual areas. To probe the functional capabilities of these representations and test existing specialization accounts implicating FFA in face identification [5], RSC in spatial cognition [47] and VWFA and EBA in letter and body perception respectively, we simulated these fine-grained discrimination tasks using the representations from response-optimized models of all ROIs. We found that the representations from the FFA network outperformed the other representations and a representation from a task-optimized network at the face identification task. We also found that the RSA network beat the other representations and a representation from a task-optimized network at the spatial task. Similarly, among response-optimized models, VWFA and EBA representations surpassed others at the letter classification and body pose estimation tasks respectively. This selective transfer to different visual discrimination tasks provides strong clues into the functional roles subserved by different representations. Moreover, the observation that these complex visual capacities are realized spontaneously in neural networks optimized solely to predict brain activity suggests that the utility of these networks may not be restricted to modeling cortical activity, rather they may serve as useful models of biological function as well.

### A possible complete characterization of selectivity in the ventral visual cortex?

When considering a mask of the ventral visual regions and training a new model to predict the entire region, we recovered the categories detected when considering the ROIs overlapping with this mask: heads (faces), signboards (words) and skin (bodies). We also obtained selectivity for food, which was also reported in recent works [37, 38, 39]. However, we did not find detectors for the thousands of other semantic concepts present in the large stimulus set and the dissection concept dictionary. This result presents an intriguing hypothesis that we have already identified all the major categories for which the ventral visual cortex is selective. It should be specified that this result might be constrained by the diversity and coverage of images in the NSD dataset, the precision of the pixel-level labels in the datasets used for network dissection, the limited spatial resolution of fMRI, by our modeling approach, and by the fact that we perform the dissection at a single voxel level. Since the NSD dataset offers high-quality response measurements afforded by the use of ultra-high magnetic field strength (7T), is already quite expansive and diverse in the stimulus set, and since the datasets used for dissection have been meticulously labeled down to the pixel-level against a broad range of concepts, it is likely that this hypothesis holds merit. This hypothesis relies on the definition of selectivity as the most activating category for a voxel or a region. As we saw in this paper, regions in the high level visual system also respond to characteristic patterns in the absence of the preferred categories. It is possible that these graded responses to non-preferred categories might still be used by upstream regions when processing non-preferred categories.

### Extensions

Probing computational models of the brain imposes several challenges that may confound our conceptions about neural representations and computations. One key issue is that any conceptual insights gained from a computational model are helpful insofar as the model is a good approximation of the biological system. The prediction accuracy of response-optimized DNN is not yet perfect. This may be due to limitations in the model architecture, insufficient stimulus-response data for fitting complex neural network models, supervision from noisy fMRI signals, or a combination of these and other factors. The work presented in this study, nevertheless, was able to 1) replicate decades of a significant body of hypothesis-driven work, 2) show the robustness of the learned representations even in the absence of the category of interest, 3) show that these representations could achieve important functional roles hypothesized to be characteristic of their respective ROIs and 4) that the category selectivities can be recovered even when the analysis approach is applied on the entire ventral visual pathway. This work also revealed a new empirical space to improve predictions of brain activity in future work, by considering different model architectures and focusing on high-order cortical regions (particularly along the dorsal visual pathway where the current task-optimized models have not yielded the same level of predictive success), or other imaging techniques (MEG, EEG, etc.).

An important aspect of the network dissection used here is that it studies units in isolation. In several cases, semantic concepts may be encoded by a combination of multiple units (voxels). Future studies could extend network dissection techniques to understand the properties of simulated population responses in underexplored regions of the visual cortex, whose precise functional characterization remains elusive. Even though single neurons or single voxels may not exhibit high selectivity for object categories in those areas of the brain, we can ask whether populations of neurons or voxels in these regions encode and represent human-understandable concepts using novel network interpretability techniques [74].

Our work demonstrates that a less hypothesis-committed approach can complement hypothesis-driven studies of the visual cortex in meaningful ways. This empirical approach, enabled by advances in data science, and large-scale compilation and dissemination of neural data, can offer a complementary basis for building broader theories about neural computations, that generalize to a range of ethologically relevant scenarios.

## Methods

### Natural Scenes Dataset

A detailed description of the Natural Scenes Dataset (NSD; http://naturalscenesdataset.org) is provided elsewhere [32]. Here, we briefly summarize the data acquisition and preprocessing steps. The NSD dataset contains measurements of fMRI responses from 8 participants who each viewed 9,000–10,000 distinct color natural scenes (22,000–30,000 trials) over the course of 30–40 scan sessions. Scanning was conducted at 7T using whole-brain gradient-echo EPI at 1.8-mm resolution and 1.6-s repetition time. Images were taken from the Microsoft Common Objects in Context (COCO) database [75], square cropped, and presented at a size of 8.4° x 8.4°. A special set of 1,000 images were shared across subjects; the remaining images were mutually exclusive across subjects. Images were presented for 3 s with 1-s gaps in between images. Subjects fixated centrally and performed a long-term continuous recognition task on the images. Informed consent was obtained from the participants and the study was approved by the University of Minnesota Institutional Review Board.

The fMRI data were pre-processed by performing one temporal interpolation (to correct for slice time differences) and one spatial interpolation (to correct for head motion). A general linear model in which the HRF is estimated for each voxel and the GLMdenoise technique is used for denoising was then used to estimate single-trial beta weights. Betas provided in the subject-native space (func1pt8mm) were used in all of our experiments. Every stimulus considered in this study had 3 repetitions. We averaged single-trial betas after z-scoring every voxel within each scan session to create our voxel responses. The 4 ROIs considered in this study, namely, the Fusiform face area (FFA, includes FFA1 and FFA2), Extrastriate body area (EBA), Visual word form area (VWFA) and Retrosplenial cortex (RSC), were manually drawn based on the results of the functional localizer (floc) experiment after a liberal thresholding procedure.

### Response-optimized encoding model

We trained separate voxel-level predictive models for each of the above category-selective regions with the same backbone architecture. The predictive model comprises a shared convolutional neural network *core* common across all subjects that represents the feature space unique for specific visual areas. We employ a linear *readout* model on top of the feature space to predict the responses of individual voxels in a specific region of interest under the assumption that the feature space likely represents the input received by these areas and these regions perform close-to-linear transformations on this input. A linear readout on a shared feature space is further based upon the often made assumption that the activity across a set of neurons or voxels in one individual can be related to the activity of the second individual in the homologous functional region by a linear transform [76]. Further, the linear readout is also *factorized* into *spatial* and *feature* dimensions following popular methods for neural system identification. This allows us to separate spatial tuning or receptive field locations (i.e., what portion of the sensory space is the voxel most sensitive to?) from feature tuning (i.e. what features of the visual input is the voxel sensitive to?).

The base feature extraction network or the core thus performs all nonlinear transformations to convert the raw sensory stimuli (i.e., pixels) into a representation characteristic of a particular visual area, whereas the readout linearly maps this extracted representation into voxel responses. The core consists of four sequential convolutional blocks, with each block comprising the following feedforward computations: two convolutional layers each followed by an inner batch norm and nonlinear activation (ReLU) operations and an anti-aliased AvgPool operation at the end. Instead of regular convolutions, we employ E(2)-steerable convolutions in the core of all our models to compute orientation dependent activations for many different orientations, thereby achieving joint equivariance under translations and rotations by design [77, 78, 43]. This enables us to apply filters not just in every spatial location, as in a standard convolutional layer, but also in every orientation, increasing parameter sharing and improving the statistical efficiency of deep learning. This modeling choice is also inspired by neural computations in early visual areas where it is hypothesized that groups of neurons perform similar computations at different orientations, e.g., edge or curve detection at different orientations. We further evaluate the usefulness of our design choices in Appendix D and Supplementary Fig. S5.

### Architectural details

Our core consists of four convolutional blocks, each comprising two convolutional layers. Each convolutional layer contains 48 feature sets where each feature is extracted at 8 orientations, resulting in 384 (48 × 8) feature maps. The core is thus equivariant under rotations by 45 degrees. Naive filter rotations can result in aliasing artifacts. From an implementation perspective, the filters in these equivariant convolutional operations are constructed as a linear combination of a fixed system of atomic filters, which helps avoid artifacts and enables arbitrary angular resolution with respect to sampled filter rotation [77]. The first convolutional block contains layers with a filter size of 5 while all the remaining layers have a filter size of 3. Each convolutional layer is followed by a BatchNorm and ReLU operation. Each block is followed by an Average Pooling operation with a stride of 2 (except for the last Average Pooling where we use a stride of 1). For Average Pooling and BatchNorm, we use the implementations provided for steerable convolutional neural networks so that the feature fields are processed in a way that ensures equivariance [78].

The readout contains all voxel-specific parameters and maps the representation extracted from the core (with dimensions 384 × 28 × 28) to individual voxel responses. Weights of the readout are a sum of outer products between a spatial filter (of dimensions 28 28 per voxel) and a feature vector (a 384-D vector per voxel). The spatial filter further had a positivity constraint (enforced using rectification) and was normalized independently for each voxel by dividing each spatial weight by the square-root of the sum of squared spatial weights across all locations.

## Training and testing models

### Input preprocessing

All input images were scaled to 224 × 224 pixels for computational efficiency. Further, all images were normalized according to the ImageNet mean ([0.485, 0.456, 0.406]) and standard deviation ([0.229, 0.224, 0.225]) for both response-optimized and task-optimized models.

### Model optimization

To develop the model, we use the first 4 of 8 subjects. Among those 4 subjects, the dataset comprises 37,000 natural scene images, among which 1,000 images are shared across all subjects and the rest are exclusive to each subject. We used the 1,000 shared images for testing our models and split the remaining stimulus set into 35,000 training and 2,000 validation images.

All parameters of the response-optimized model were optimized jointly to minimize the *mean squared error* between the predicted and measured response. Since for every image in the training set, the response is measured from only a single subject and not all subjects, we use a masked mean squared loss to train the model across multiple subjects. Let 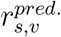 and 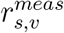. denote the predicted and measured response of voxel *v* in subject *s* to image *i*, respectively. Then, the loss function during training is given, as

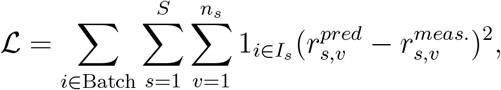

where 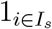 is the indicator variable specifying if image *i* was shown to subject *s*. The proposed method allows us to propagate errors through the shared network even if the subjects are not exposed to common stimuli since we can always exclude the subjects/voxels for which the response is not present from mean error calculation within each batch. The shared network thus benefits from diverse, varying stimuli across subjects with less extensive constraints on data collection from single subjects. Models were trained for a maximum of 100 epochs using Adam with a learning rate of 1e-4, a batch size of 16 and early stopping (patience = 20) based on the Pearson’s correlation coefficient between the predicted and measures responses on the validation set; validation curves were monitored to ensure convergence.

We measure performance (‘predictive accuracy’) on 1,000 test images by computing the Pearson’s correlation coefficient between the predicted and measured fMRI response at each voxel.

## Baseline models

### Task-optimized models

We compared response-optimized models against standard task-optimized models which have shown state-of-the-art performance in predicting neural responses in the primate visual cortex. In all comparisons, we employed an AlexNet architecture [79] optimized for object recognition on the large-scale ImageNet dataset [80]. We extracted features from intermediate layers of this network and employed the same spatial x feature factorized readout as used in the response-optimized networks to linearly map layer activations to brain voxel responses in each region. We selected the model layer that maximally predicted the brain responses in each region on a validation set(Conv-5 for all considered high-level visual areas). The readout parameters for task-optimized models were optimized independently for each visual region using the same training protocol as the response-optimized models. Thus, the readout models were trained for a maximum of 100 epochs using Adam with a learning rate of 1e-4 and a batch size of 16. We further applied an early stopping criterion (patience = 20) based on the Pearson’s correlation coefficient between the predicted and measures responses on the validation set.

### Categorical models

Category ideal observer models employ the category membership of labeled objects in the image to predict the responses evoked by the image. Unlike task-optimized and the proposed response-optimized models, categorical models are not image-computable and rely on annotations generate by human observers. These oracle models have absolute access to the categories present in an image and have previously been shown to explain substantial variance in image representations in both macaque and human IT [81, 21]. One might expect their performance to be even higher for explaining image representations in category-selective visual clusters in high-level cortex. We obtained object category labels for every NSD image from the MS COCO database [75]. The input to the categorical model is thus an 80-D binary vector corresponding to the 80 object categories annotated in the database, where each element indicates whether the corresponding category was present in the image or absent (note that NSD images contain multiple objects per image). We fitted *l*_2_ regularized linear regression models (known as ridge regression) on this representational space to find weights corresponding to different categories for every voxel. The regularization parameter was optimized independently for each subject and for voxels in each visual area by testing among 8 log-spaced values in [1e-4, 1e4].

### Generalization to new subjects

We tested the generalization of different competing models to a set of 4 new subjects in the Natural Scenes Dataset by freezing the weights of these models and only training linear readouts on top of their fixed representations. The readouts had the same architecture as the original models, i.e., they were factorized into spatial and feature dimensions. The spatial map had a positivity constraint and was again normalized independently for each voxel by dividing each spatial weight by the square-root of the sum of squared spatial weights across all locations. The linear readouts were trained with a gradient descent procedure (Adam optimizer, learning rate 1-4) that minimizes the mean squared error between the voxel-wise predictions and true measured responses. For each new subject, the dataset comprised brain responses for 5,445 stimuli with 3 repetitions (remaining stimuli had either 1 or 2 repetitions and were discarded from analysis). We varied the size of the training set from a mere 100 stimulus-response pairs to a large set of 4,500 pairs. The readouts were trained independently for each subject, each training set size and each visual ROI for 100 epochs with an early stopping criterion (patience of 20). We monitor the loss on a validation set comprising 430 samples. The performance is evaluated on an independent set of 515 stimulus-response pairs shared across the new subjects using Pearson’s correlation coefficient (R). These stimuli were not encountered during the training of response-optimized models on the original set of 4 subjects. An identical readout training procedure was followed for both response-optimized and task-optimized models. In the case of categorical models, we follow the same procedure that we applied on the original set of subjects and fit *l*_2_ regularized linear regression models on the categorical representational space to find weights corresponding to different categories for every voxel. The regularization parameter was optimized independently for each subject, each training set size and for voxels in each visual area by testing among 8 log-spaced values in [1e-4, 1e4].

### Quantifying the semantic selectivity of voxels

High-level visual concepts are generally verbalizable, and throughout this paper, we refer to voxels that encode and represent these concepts as ‘semantically selective’. To quantify the selectivity of voxels for different human-interpretable concept categories, we adapt the previously proposed framework of ‘Network dissection’ [35, 36] to our brain response-optimized models. We see this as a fine-grained approach to characterize voxels that looks at not just the image-level category labels but rather dense pixel-level segmentations across thousands of cluttered natural scenes to characterize a voxel. The probe dataset used for quantifying the semantic selectivity of voxels comprises 36,500 held-out images from the validation set of the large-scale Places365 dataset. The reference segmentation for these probe images comes from the Unified Perceptual Parsing image segmentation network [82] previously trained on 20,000 scene-centric images from the ADE20k dataset [83]. The ADE20k dataset is exhaustively and densely annotated with objects, parts of objects and in some cases, even parts of parts. This reference segmentation assigns every pixel a semantic label from a large vocabulary of human-interpretable concepts, comprising 335 object classes, 1,452 object parts and 25 materials.

A unique advantage of the factorized readout employed in this study is that it allows us to disentangle spatial selectivity from feature selectivity. To enable the network dissection procedure to be applicable to our models, we first discard the learned spatial selectivity of every voxel and use the learned feature tuning of every voxel to create an additional 1×1 convolutional layer, so that every voxel is represented by an independent unit in this convolutional layer. These units are used to characterize the semantic selectivities of voxels irrespective of the position of the respective semantic categories in the visual field. This yields a model that is entirely convolutional. Dissecting the last layer of this model (which has as many units as the number of voxels in the ROI) reveals the semantic selectivities of all voxels. We only performed the dissection procedure on voxels that had a held-out raw prediction accuracy (R)*>*0.1 and the total number of such voxels in each visual ROI across 4 subjects are listed in Table 1. As proposed in [35], the selectivity of a particular unit (or voxel in our case) is quantified by computing the Intersection over Union (IoU) of the corresponding thresholded and binarized activations of that unit for a large number of images from the probe dataset against the reference segmentation. This metric computes the ratio of the number of pixels common between both spatial maps and the total number of pixels activated in either the reference segmentation map or the binarized activation map. A voxel is termed as semantically selectivity for a *concept* if its IoU with the reference segmentation of that concept is greater than 0.04. Further details about the dissection procedure employed in this study are described elsewhere [36]. Our main models are trained on the voxels of one ROI at a time. We evaluate the effect of using a shared model across ROIs on network dissection in Appendix B and the effect of fitting the single ROI models to new subjects in Appendix C.

### Synthesizing maximally exciting inputs

We performed a qualitative *feature visualization* analysis to find the visual pattern that would maximally activate individual model neurons emulating brain voxels. Neural networks are differentiable with respect to their inputs. Starting from a random noise input, we use these gradients to iteratively move the input towards the goal of maximizing activation in individual model neurons. This visualization technique is commonly employed in neural network interpretability research to find the features that drive model neurons [84]. Most visualization techniques further employ an *image prior* in the form of a regulariser to restrict the maximally exciting input to a suitable subset of the image space [85, 86]. Formally, the goal of finding the maximally exciting input (MEI) *x*^*\**^ is then expressed as the following optimization problem.

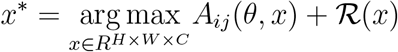

where *A*_(*i,j*)_(*θ, x*) denotes the activation of unit *i* from layer *j* in the neural network to input *x* (H: Height, W: Width, C: Channels), and *θ* denotes the parameters of the network. The latter are fixed during the above optimization procedure. *ℛ* (*x*) denotes the regulariser. In order to generate MEI for the *j*th voxel, we set *i* to the network output layer and *j* to be the index of the model neuron in the output layer that emulates voxel *j*. The above optimization problem is, in general, a non-convex optimization problem but we can find (at the very least) a local minimum by performing gradient ascent in the input space and updating *x* iteratively in the direction of the gradient of *A*_*ij*_(*θ, x*) + *ℛ* (*x*). We employed a general weak form of regularization, called ‘transformation robustness‘[87], where we stochastically jitter (up to 8 px), rotate (up to 10 degrees) and scale (by a factor between 0.8 to 1.2) the image before each optimization step so that the optimized images don’t contain high frequency noise. The images are optimized starting from random noise with Adam optimizer for 512 steps using a learning rate of 1e-3.

### Visual diet experiments

In the first set of visual deprivation experiments, we removed all stimuli belonging to the ‘person’ category (containing faces as as well as bodies). This leaves much fewer 17,325 stimulus-response pairs for training, as compared to the original 35,000 pairs that were used for training response-optimized models in the main experiment. We trained response-optimized models for two visual ROIs, namely FFA and EBA, with the same architecture and training procedure as employed for the original models, but only changed the visual diet to the aforementioned deprived set. In another set of experimental results, shown in Supplementary Fig. S6, we removed all stimuli containing animate things, finally resulting in 12,559 stimulus-response pairs and trained another set of models for FFA and EBA on this inanimate subset. These selective reductions in training set size and characteristics resulted in a small drop in the overall prediction accuracy on the complete test set, as reported in Supplementary Fig. S9, but did not result in selective disruptions for the out-of-domain category versus in-domain stimuli.

### Transfer learning on fine-grained visual discrimination tasks

To formally test existing functional specialization accounts which implicate FFA in face perception, RSC in spatial cognition, VWFA and EBA in letter and body perception respectively, we simulated several fine-grained discrimination tasks with independent stimuli in response-optimized models of all brain regions, as mentioned below:

- *Face identity discrimination task:* This includes stimuli from the MiniCelebA dataset which comprises facial images of 20 identities, each having 100/30/30 train/validation/test images [58].
- *Spatial layout estimation task:* Here, the stimuli include 4,000 diverse indoor scenes from the SUN database [63]. These stimuli were split into sets of 3200 training, 400 validation and 400 test images. Each stimulus image has a corresponding label for the room layout type, where the layout categories were defined using a keypoint-based parameterization (as illustrated in Figure 7). This helps us frame the room layout estimation task as a classification problem.
- *Letter classification:* The printed letters dataset, generated in prior work [64], contains 65,520 images of the 26 uppercase Latin letters. Here, the same letter appears in different fonts (out of fourteen), different sizes (out of five), different weights (bold or not bold), different styles (italicized or not italicized) and different pixel locations (out of nine), creating diverse visual appearances. To make the classification task more difficult, we used a maximum of 1,000 images across all letters (split into 80%training and 20% validation) for training and reserved the remaining 64,520 images for testing.
- *Body pose classification:* Here, the dataset includes images of multiple computer-graphics (CG)-generated models doing 5 different yoga poses [65], divided into 1,006 (split into 80%training and 20% validation) and 501 test images. The images are rendered on multiple different backgrounds, and contain both male and female models with a variety of skin and hair tones, captured from different viewpoints and lighting.

We use response-optimized models to extract the *predicted* responses of voxels in every region to stimuli from the fine-grained visual categorization tasks. To ensure that the differences in performance of response-optimized models on fine-grained visual categorization are not driven by the differences in the number of voxels in every region, we selected the top 512 voxels in every region based on test correlations (i.e., correlation between the predicted and measured responses on 1,000 test images). This number was chosen to match the dimensionality of representations from the pre-trained VGG16 architecture [60] optimized for face-recognition on the large-scale VGGFace2 dataset [88]. We consider the performance of the VGG16 representation on the MiniCelebA dataset as an estimate of the upper bound on performance expected by our models on face recognition (we did not have access to human face recognition performance on this dataset). We also consider the 512-D dimensional representation from a VGG16 architecture trained on image categorization using the large-scale ImageNet database as an additional baseline for all visual discrimination tasks. This baseline was chosen because ImageNet-trained networks yield highly transferable representations that perform well in a range of vision-based tasks [61].

We fitted *l*_2_ regularized linear classification models (known as ridge classifiers) on these different representational spaces to predict the class label (e.g. facial identity or room layout type) of held-out stimuli. We vary the size of each transfer dataset from 60-100% of the maximum training set size and report the corresponding classification accuracy on the held-out set as a function of the transfer dataset size. For each of the above representational models (response-optimized, ImageNet-optimized or VGGFace2-optimized), the regularization parameter was optimized independently for each task (face recognition, spatial layout estimation, letter classification and body pose classification) and each training set size (60-100%) by testing among 10 log-spaced values in [1e-5, 1e5]. We selected the regularization parameter value that yielded best classification accuracy on the validation dataset.

### Training a model on the entire ventral visual cortex

To test whether category selectivities may be uncovered with an impartial approach that does not require ROI selection or localization, we trained a single response-optimized model on responses from the entire ventral visual stream across all subjects. The model employed the same architecture as the ROI-specific models and was trained following the same procedure as employed for ROI-specific models and described above. The ventral visual stream voxels in every subject were extracted using the streams atlas released with the Natural Scenes Dataset [32]. Briefly, this ROI collection reflects large-scale divisions of the visual cortex into primary visual cortex and intermediate and high-level ventral, lateral and dorsal visual areas. These were manually drawn by NSD curators for each subject after accounting for the voxel-level reliability metrics. We extracted the ROI mask corresponding to the higher-level ventral stream label. This ROI was drawn to follow the anterior lingual sulcus (ALS), including the anterior lingual gyrus (ALG) on its inferior border and to follow the inferior lip of the inferior temporal sulcus (ITS) on its superior border. The anterior border was drawn based on the midpoint of the occipital temporal sulcus (OTS). As shown in Figure 8, it is very broad (7,000-9,000 voxels per subject), yielding a total of 28,910 voxels across all 4 NSD subjects.

### Noise ceiling estimation

Imperfect predictions of models are not solely due to model limitations, but may arise due to the inherent noise in the fMRI signal, which biases the prediction accuracy downward. Noise ceiling for every voxel represents the performance of the “true” model underlying the generation of the responses (the best achievable accuracy) given the noise in the measurements. They were computed using the standard procedure followed in [32] by considering the variability in voxel responses across repeat scans. The dataset contains 3 different responses to each stimulus image for every voxel. In the estimation framework, the variance of the responses, 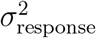, are split into two components, the measurement noise 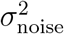 and the variability between images of the noise free responses 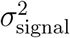.

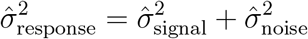

An estimate of the variability of the noise is given as 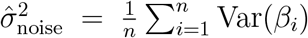, where i denotes the image (among *n* images) and Var(*β*_*i*_) denotes the variance of the response across repetitions of the same image. An estimate of the variability of the noise free signal is then given as,

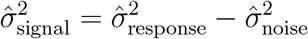

Since the measured responses were z-scored, 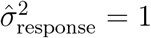 and 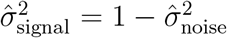. The noise ceiling (n.c.) expressed in correlation units is thus given as 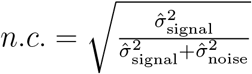. The models were evaluated in terms of their ability to explain the average response across 3 trials (i.e., repetitions) of the stimulus. To account for this trial averaging, the noise ceiling is expressed as 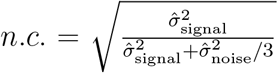. We computed noise ceiling using this formulation for every voxel in each subject and expressed the noise-normalized prediction accuracy (R) as a percentage of this noise ceiling.

## Supporting information

Supplementary Results

## Acknowledgements

The authors thank the Natural Scenes Dataset (NSD) team for collecting and sharing the dataset. This effort was led by PIs Kendrick Kay and Thomas Naselaris, and includes contribution by Emily J Allen, Ghislain St-Yves, Yihan Wu, Jesse L Breedlove, Jacob S Prince, Logan T Dowle, Matthias Nau, Brad Caron, Franco Pestilli, Ian Charest and J Benjamin Hutchinson.

## Author contributions

M.K. and L.W. conceptualized the research. M.K. built the model and ran the analyses.

M.K. and L.W. wrote the paper.

## Code

The code used in this paper can be found at https://drive.google.com/drive/folders/1YJqLNFE1fwuHSIVGC_ZIxos87jUSNI8X?usp=sharing. It will be released as a Github repository upon acceptance.

## Data

We use the open 7T fMRI Natural Scenes Dataset (NSD) [32].

## Footnotes

1 Pre-trained model downloaded from here: https://pytorch.org/hub/pytorch_vision_alexnet/

