## Supplementary Results for "Higher visual areas act like domain-general filters with strong selectivity and functional specialization"

### Supplementary analysis

**Appendix A - Face selective units in models of EBA and VWFA.** The network dissection procedure shown in Fig. 2 revealed face-selective units in models of EBA and VWFA. While these ROIs showed some overlap with FFA due to the liberal thresholding procedure (so the presence of face detectors in these models was not surprising), we asked whether these face-detecting units were indeed selective for faces in the independent functional localizer experiment. We thus compared the IoU of units, as quantified by the dissection procedure, against the t-value index of face selectivity, as quantified by the localizer. As shown in Fig. 5 and Supplementary Fig. S1, we get a reasonably high match between the two quantities, despite the vast differences in the experimental paradigm and quantification procedure involved in the two. The high correlations suggest that the model can tease apart face-selective voxels among a group of voxels selective for another visual category and, more generally, can meaningfully capture fine-grained distinctions between voxels within the same visual region.

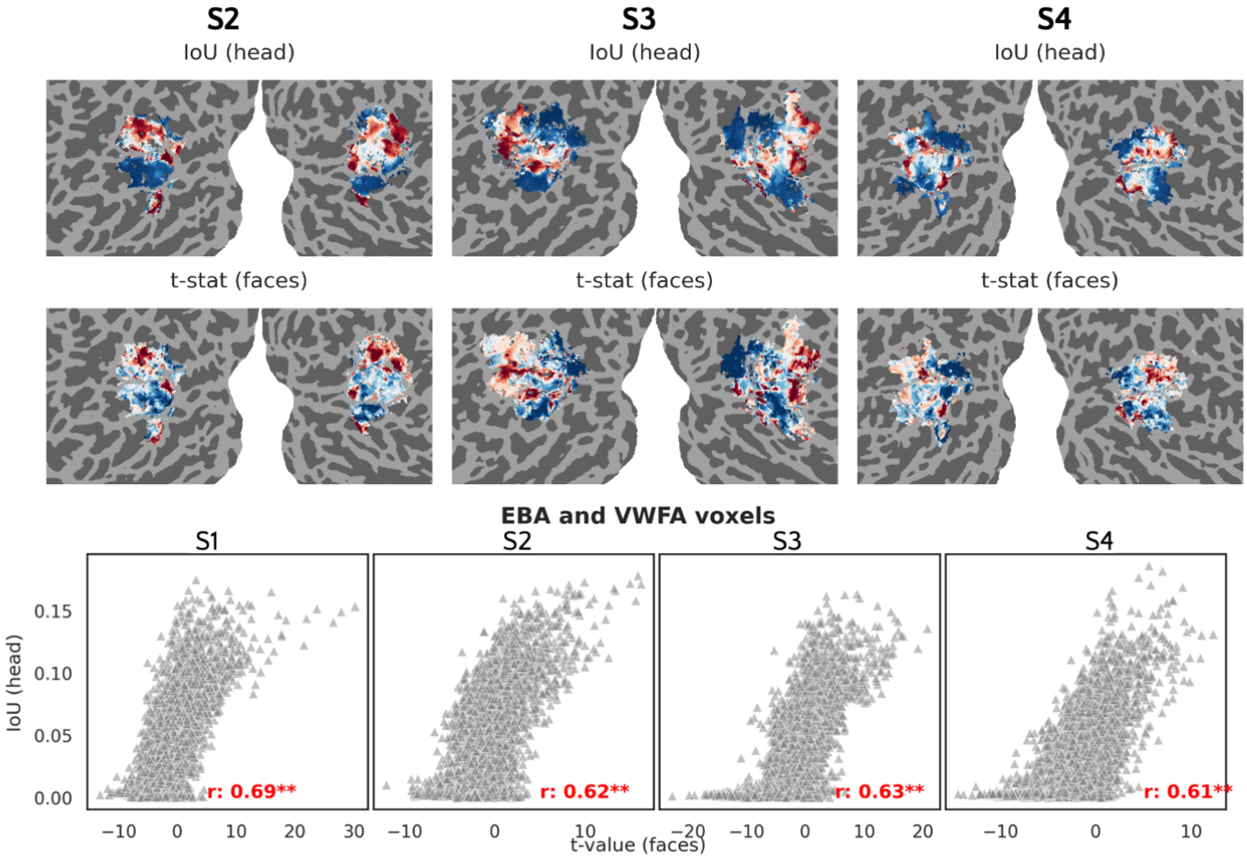


**Supplementary Figure S1. IoU versus t-statistic for faces.** [Top] Cortical surface plot of face selectivity as measured by the localizer experiment against the IoU with the 'head' concept as quantified by the dissection procedure for the remaining three subjects (subject 1 is shown in Figure S2). [Bottom] Scatter plot of IoI with 'head' against face-selectivity measured with the independent localizer (t-value) for all EBA and VWFA voxels.

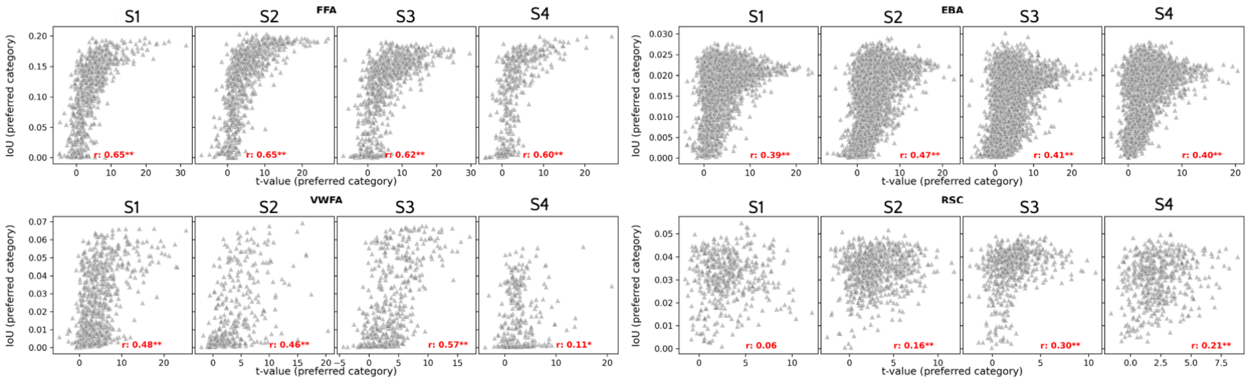


**Supplementary Figure S2. Within-ROI variability in voxel responses**. Agreement between semantic alignment of individual voxels measured with the network dissection procedure and their category-selectivity measured with the independent functional localizer experiment. Each point is an individual voxel within the ROI: y-axis indicates its IoU with the preferred concept of the respective ROI ('head', 'signboard', 'person' and 'window' for FFA, VWFA, EBA and RSC, respectively) and x-axis represents the t-value index of selectivity against the preferred category in the localizer ('faces', 'words', 'bodies' and 'places' for FFA, VWFA, EBA and RSC, respectively).

**Appendix B - Network dissection of combined model.** As described in the main text, we modeled the responses of the 4 visual ROIs considered in the study separately. These regions were identified with an independent functional localizer. However, this approach still required prior localization of these category-selective regions and is biased by the hypothesis that these regions perform distinct computations. We wanted to assess whether semantically selective units would emerge in a model trained to predict the responses of all voxels simultaneously. In theory, this is a less hypothesis-committed approach as we don't commit to a set of voxels and let the data-driven method 'discover' the semantically selective units among them. We trained a joint model for predicting the responses of all voxels in the ventral visual stream (totalling 28,910 voxels across all 4 subjects). This method helps us answer (a) whether a combined model performs better or worse than ROI-specific models and (b) whether it is capable of learning highly varying semantic concepts all at once and distinguishing between the high-level selectivity of voxels in each region. As shown in Figure S7, we again found strong evidence of category selectivity, aligned with the hypothesized functional role of these ROIs. Further, the combined model performs just as well as ROI-specific models, as shown in Figure S7A, indicating that a single model is sufficient to account for diverse tuning properties throughout the ventral visual stream and we are investigating this model further in future work.

**Appendix C - Network dissection on ROI-specific response-optimized models of new subjects.** In Figure 1E, we showed that response-optimized models generalize to novel subjects with limited samples. In fitting these models to new subjects, we only trained the linear readout while keeping the rest of the network fixed. We asked whether units in these tuned models also specialized in recognizing highly specific semantic features of inputs. We

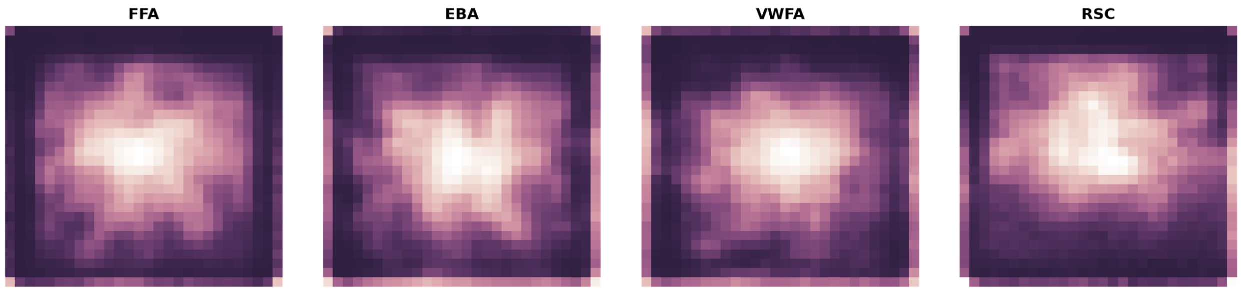


**Supplementary Figure S3. Spatial masks**. The learned spatial weights of all voxels in the response-optimized model are averaged to create an ROI-specific spatial mask. The spatial masks have a large spread around central vision, consistent with the known spatial receptive field properties of these higher-order ROIs. RSC appears to show an upper visual field bias.

| ROI | Number of voxels dissected |
|---|---|
| FFA | 2,721 |
| VWFA | 1,840 |
| EBA | 12,732 |
| RSC | 2,336 |

Table 1: Number of dissected units in each model, i.e. the number of voxels that had raw prediction accuracy (R) greater than 0.1

dissected these models to quantify how the units (i.e., model voxels) responded to individual concept categories like colors, textures, objects and parts of objects. Supplementary Fig. S4 shows the results of this analysis. Similar to results we reported in the main text, we again see highly specific semantic concepts aligned with hypothesized functional role of different ROIs and a similar pattern of selectivity across visual ROIs.

**Appendix D - Evaluating architectural choices and training schemes.** Training response-optimized models requires a large number of stimulus-response pairs to achieve an accuracy comparable to task-optimized models trained with millions of images. We found that architectural design choices had a considerable impact on the performance of response-optimized models. In particular, sharing weights across filter orientations and utilizing a factorized readout (thereby, enforcing sparsity) both enable sample-efficient learning.

We investigated the effect of a sparse readout on the task-optimized model in a separate ablation study. Task-optimized models have a fixed core so the precise effect of different readout schemes is more dissociable in these models. We compared two conditions:
*(1) Fully connected non-sparse readout:* Here, we use a linear fully connected layer to map the features from the representational core network to voxel-level responses across the initial set of 4 subjects. Thus, if the output of the representational core has dimensions $H \times W \times C$and there are $V$ voxels, the number of parameters in this readout are $H \times W \times C \times V$. See

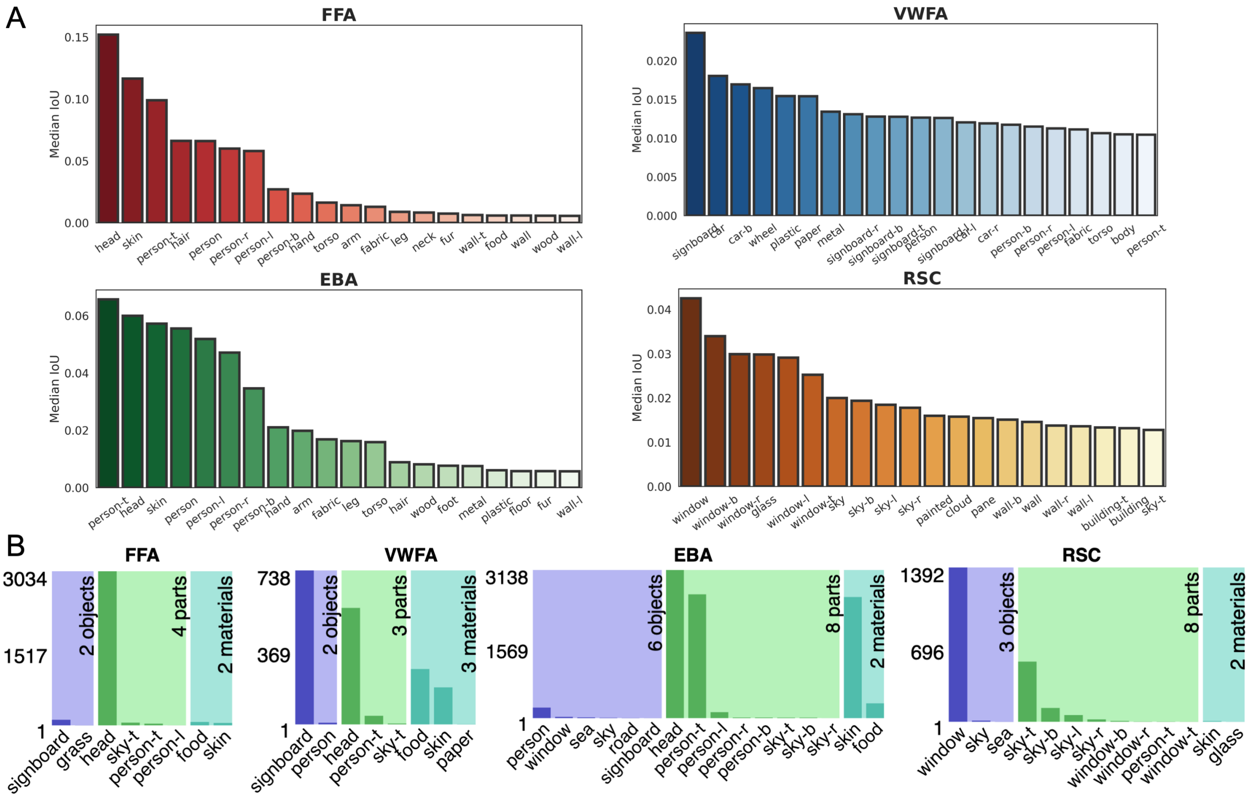


**Supplementary Figure S4. Dissection of models tuned on new subjects.** **A** demonstrates the median IoU metric across all voxels belonging to an ROI for the top 20 visual concepts rank ordered by median IoU. The top concepts for FFA, VWFA, EBA and RSC discovered using this hypothesis-neutral approach (‘head’, ‘signboard’, ‘person’ and ‘window’ respectively) align remarkably well with the known domain-specificty of voxels in these regions and with the results obtained from dissection of the original models in the main experiment. **B** shows the matched concepts for every tuned response-optimized model, i.e., the number of units in the ‘voxel’ layer that showed high alignment (stringent threshold of IoU>0.04) with a human-interpretable visual concept. Again, similar to results reported for the original models trained on the initial set of 4 subjects, the matched visual concepts in these tuned response-optimized networks are also highly specific and aligned with the previously hypothesized functional role of these ROIs.

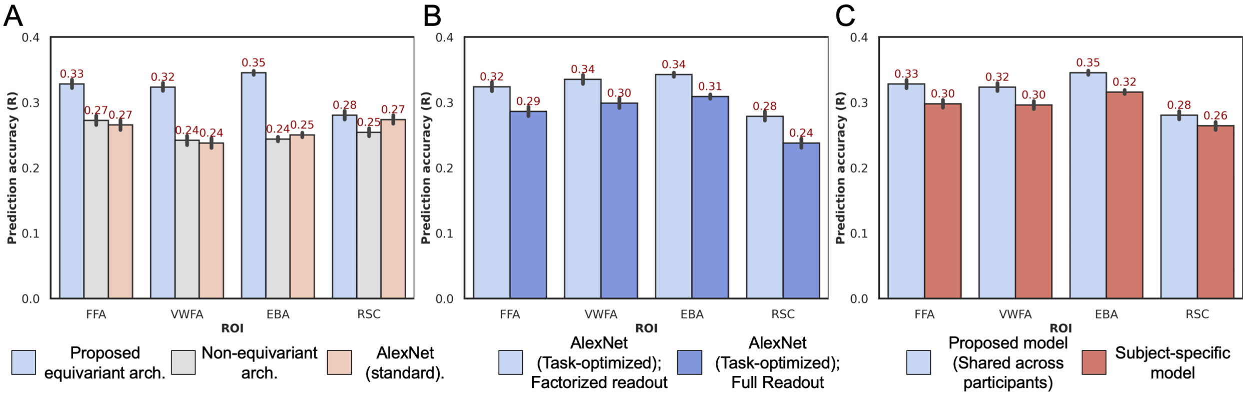


**Supplementary Figure S5.** Quantifying the effects of core/readout architectures and cross-subject training on prediction accuracy of the response-optimization approach. **A** shows the quantitative advantage of using a rotation-equivariant core over comparative non-equivariant and standard non-equivariant (AlexNet) architectures. The equivariant core consistently outperforms the latter architectures in response prediction for held-out images by a significant margin. **B** shows the quantitative advantage of using a factorized readout over a fully connected non-sparse readout method for the AlexNet model optimized on object recognition using the ImageNet database. **C** shows the quantitative advantage of sharing a common core across subjects (with a subject-specific readout) instead of optimizing the entire core + readout model with subject-specific data. The benefits are marginal, yet significant, indicating conserved neural representations of visual stimuli across subjects.

Supplementary Fig. S5A.
*(2) Factorized readout:* Here, we use the linear readout employed in all the main experiments. This readout disentangled spatial selectivity or retinotopic biases from feature selectivity. Here, the weights are a sum of outer products between a spatial filter and a feature vector and the number of parameters are $(H \times W + C) \times V$, a dramatic reduction in the number of parameters from the fully connected layer with a factor of nearly C (256 in the case of the AlexNet model). See Supplementary Fig. S5B.

Next, we investigated the effect of training a shared non-linear representation over multiple subjects instead of learning a subject-specific representation. In this ablation study, we trained response-optimized models using only subject-specific data. Note that this also reduced the number of samples from 35,000 stimulus-response pairs to 8,500 paired samples for response optimization. We train these models separately for each subject and monitor validation accuracy on a set of 500 samples. The final prediction accuracy (R) is again computed on the independent set of 1,000 images. As shown in Supplementary Fig. S5C, single-subject models perform slightly worse than models that share a convolutional core across subjects, indicating the quantitative advantage of building shared models. The fact that a shared model outperforms single-subject models further suggests that neural representations of visual stimuli are largely conserved across subjects. Further, the strong generalization of these shared convolutional cores to new subjects with the limited-samples transfer learning paradigm suggests that they capture fairly general cortical features characteristic of a visual ROI and not subject-level idiosyncrasies.

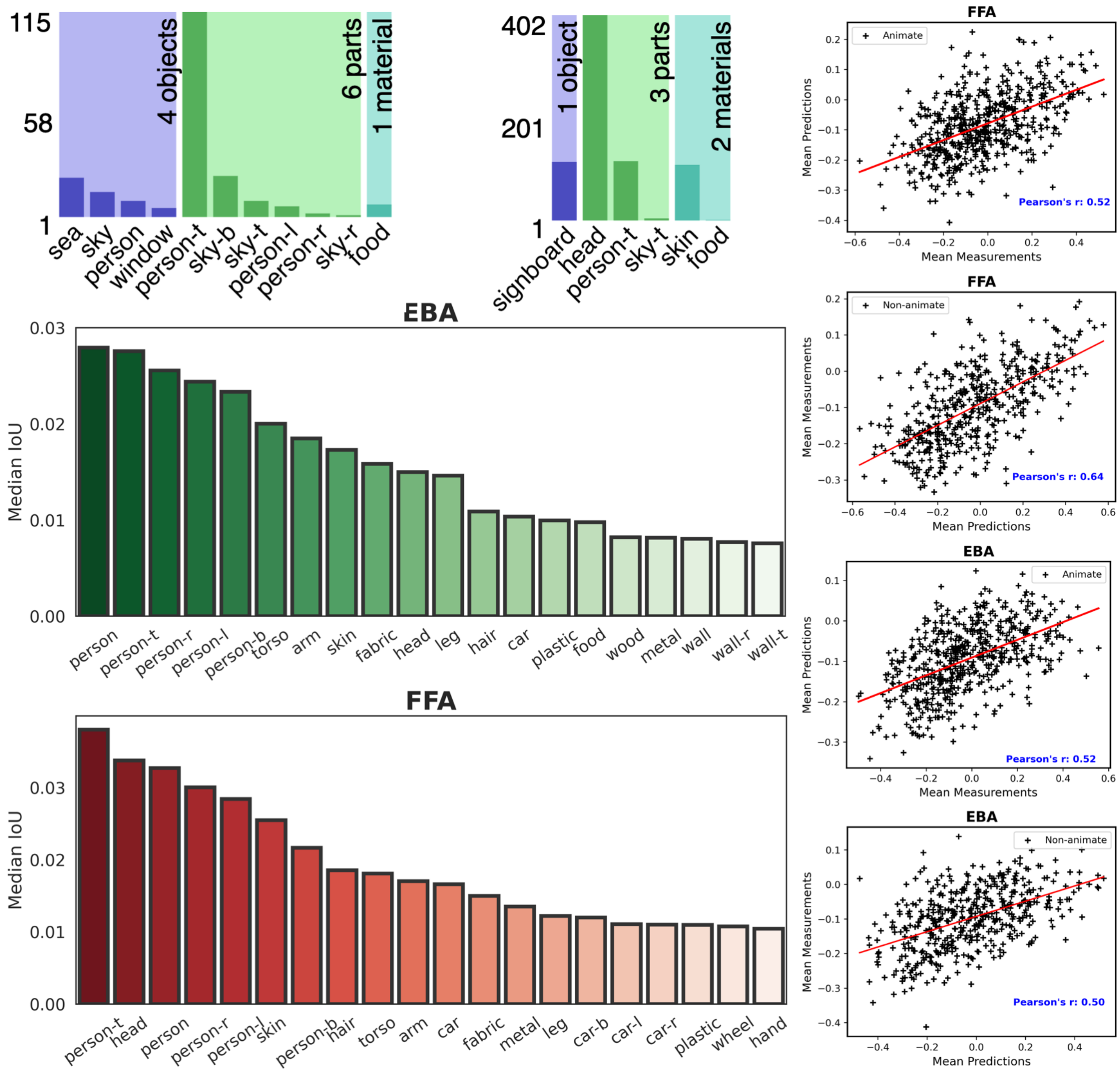


**Supplementary Figure S6. Other deprivations in visual diet**. Results of the dissection procedure applied to response-optimized models of FFA and EBA respectively, when trained using a visual diet entirely deprived of all animate categories. This leaves much fewer 12,559 stimulus-response pairs for training, as compared to the original 35,000 pairs that were used for training response-optimized models in the main experiment. Despite the extensive deprivation, these regions are still selective for their preferred concept: EBA units show highest IoU with the 'person' category and FFA units align best with 'person-t' ('t' stands for the 'top' part of person) and 'head' concepts. The deprived models generalize remarkably to images from a new domain ('animate'), incurring only a small loss in prediction accuracy for this domain. This strong generalization, despite no experience with these visual categories, suggests that they extract domain-general visual characteristics.

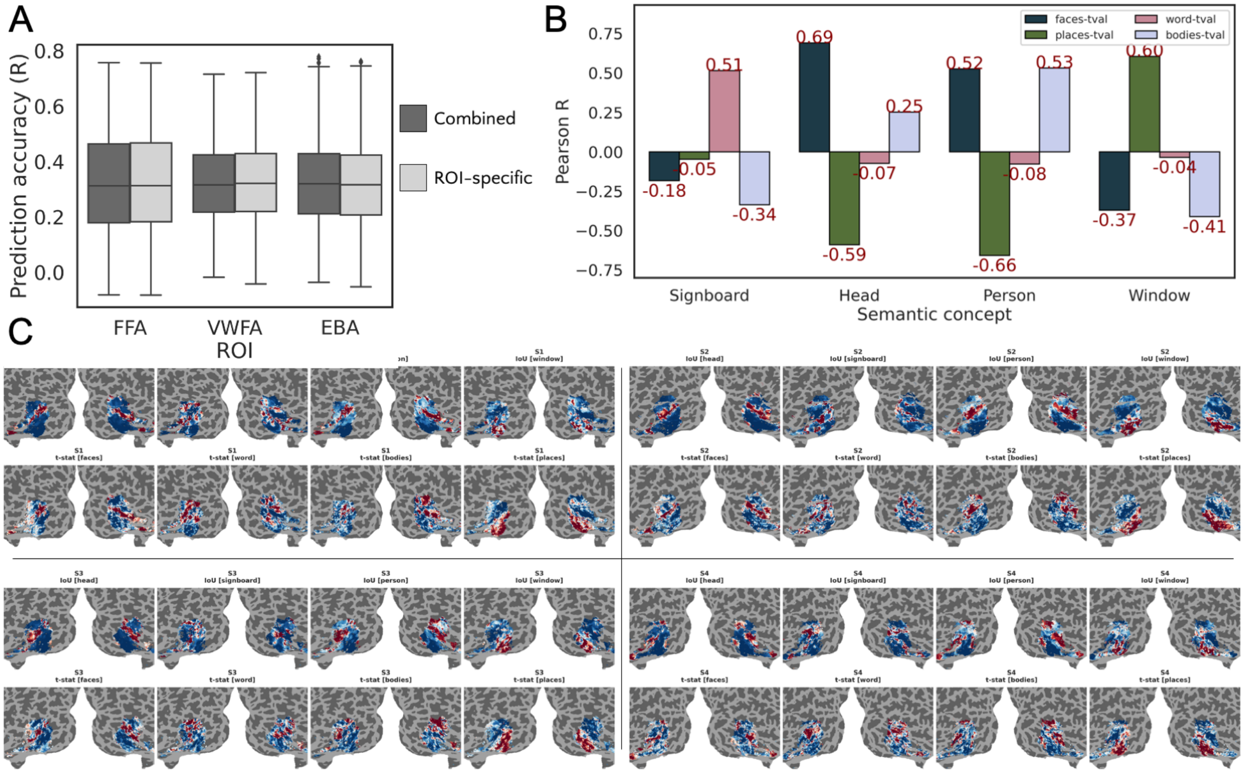


**Supplementary Figure S7.** Dissection results of the response-optimized model trained to predict brain responses across the entire ventral visual stream. The ventral ROI was drawn to follow the anterior lingual sulcus (ALS), including the anterior lingual gyrus (ALG) on its inferior border and to follow the inferior lip of the inferior temporal sulcus (ITS) on its superior border. The anterior border was drawn based on the midpoint of the occipital temporal sulcus (OTS). **A** shows the prediction accuracy of voxels that lie both within the ventral visual stream and each localized ROI (as defined previously) for two models: ROI-specific models described in the main study and a combined model that predicts the responses of all voxels in the ventral visual stream simultaneously. **B** shows the agreement between IoU of all ventral visual stream voxels against individual semantic concepts and selectivity for specific categories shown in the independent localizer experiment. Agreement is quantified by computing the Pearson's correlation coefficient between two vectors: one containing the IoU of all voxels against a concept and another vector containing t-statistic from the localizer contrast for different categories. Cortical surface plots in **C** show the IoU of every ventral visual stream voxel with the 4 visual concepts ('head', 'person', 'signboards', 'windows') that we previously found to be associated with each ROI. Below each plot, we also show the selectivity for corresponding categories (t-statistic) as computed from the independent localizer experiment; the two rows are strikingly similar (qualitatively). Each quadrant is an individual subject.

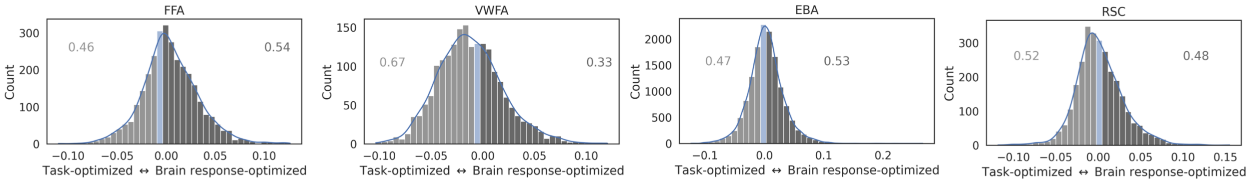


**Supplementary Figure S8. Comparative advantage plot.** Voxel-wise distribution of the difference between prediction accuracies of the response-optimized and task-optimized models for each visual ROI. The inset shows the proportion of voxels that are better predicted by each. The response-optimized models achieve parity with the task-optimized model trained on a million ImageNet images (no difference was found through a permutation test, $p>0.01$ for all 4 ROIs).

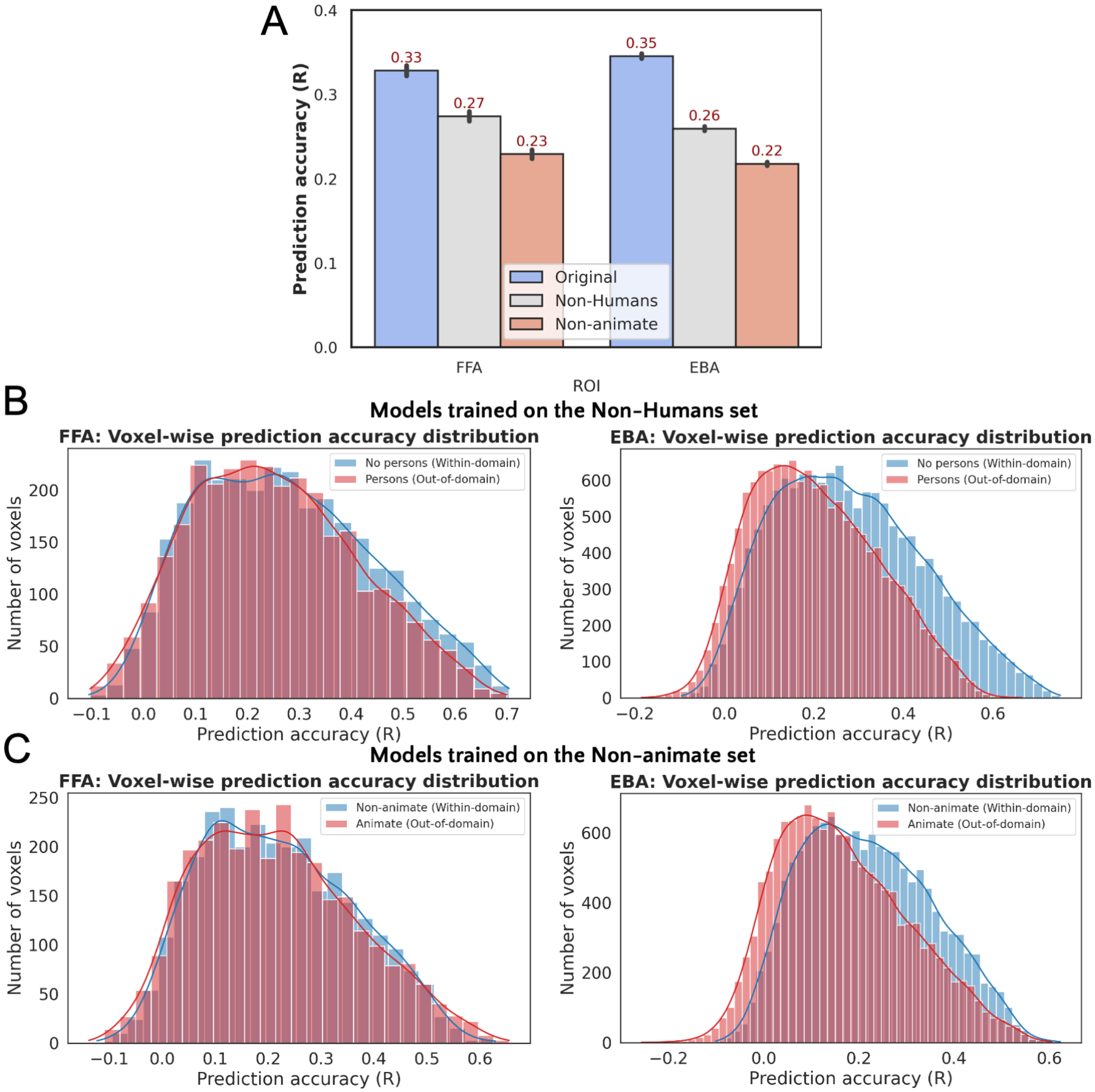


**Supplementary Figure S9. Voxel-wise prediction accuracy (R) in the deprivation experiment.** **A** shows the prediction accuracy of the FFA and EBA response-optimized models when trained using the entire dataset (Original) as well as subsets with specific deprivations (Non-Humans and Non-animate). The performance drops in the deprivation case but this drop is not surprising given the reduction in the training set size from ∼35,000 samples in the 'Original' case to 17,325 and 12,559 samples in the 'Non-Humans' and 'Non-animate' cases, respectively. Importantly, these deprivations do not result in selective disruptions for the out-of-domain category versus in-domain stimuli, and the models generalize to unseen categories remarkably well. **B** and **C** show the distribution of prediction accuracy (R) across all voxels on held-out stimuli from *within-domain* and *out-of-domain* subsets. In FFA, there is almost no performance gap between within-domain and out-of-domain stimuli for each deprivation set (**B** and **C** left). In EBA, the performance is higher for within-domain images than out-of-domain stimuli (**B** and **C** right), but the improvement is marginal.
